# Mapping RNA-binding proteins in human B cells and T cells upon differentiation

**DOI:** 10.1101/2021.06.10.447413

**Authors:** Nordin D. Zandhuis, Benoit P. Nicolet, Monika C. Wolkers

## Abstract

B cells and T cells are key players in the defence against infections and malignancies. To exert their function, B cells and T cells differentiate into effector and memory cells. Tight regulation of these differentiation processes is key to prevent their malfunction, which can result in life-threatening disease. Lymphocyte differentiation relies on the appropriate timing and dosage of regulatory molecules, and post-transcriptional gene regulation (PTR) is a key player herein. PTR includes the regulation through RNA-binding proteins (RBPs), which control the fate of RNA and its translation into proteins. To date, a comprehensive RBP expression map throughout lymphocyte differentiation is lacking. Using transcriptome and proteome analyses, we here provide an RBP expression map for human B cells and T cells. We observed that even though the overall RBP expression is conserved, the relative RBP expression is distinct between B cells and T cells. Differentiation into effector and memory cells alters the RBP expression, resulting into preferential expression of different classes of RBPs. For instance, whereas naïve T cells express high levels of translation-regulating RBPs, effector T cells preferentially express RBPs that modulate mRNA stability. Lastly, we found that cytotoxic CD8^+^ and CD4^+^ T cells express a common RBP repertoire. Combined, our study reveals a cell type-specific and differentiation-dependent RBP expression landscape in human lymphocytes, which will help unravel the role of RBPs in lymphocyte function.

## INTRODUCTION

B cells and T cells are essential to eradicate microbial infections and malignant cells. Upon antigen recognition through their receptors, B cells produce antibodies and T cells produce cytokines and chemokines, respectively. Cytotoxic T cells also acquire the capacity to kill target cells. The critical contribution of these lymphocyte subsets to anti-microbial and anti-tumor responses was evidenced by the discovery of genetic mutations in humans that result in immune dysfunction in response to infections (1). Similarly, effective T cell responses are key for tumor immunosurveillance (2).

Importantly, tight regulation of B cells and T cells effector function is key for effective clearance of infections. The aberrant production of antibodies by B cells, and the overproduction of effector molecules by T cells has been correlated with several autoimmune disorders, including systemic lupus erythematosus rheumatoid arthritis and multiple sclerosis (3–6). Likewise, patients suffering from severe disease upon COVID-19 infection developed auto-antibodies against type-I interferons (7), and an excess cytokine production in COVID-19 patients can result in organ dysfunction (8). Conversely, in chronic HIV infections or in tumors, T cells gradually lose their capacity to produce effector cytokines and to kill target cells (9,10). These findings combined highlight the necessity to fine-tune the effector function of B cells and T cells.

To perform their effector function, B cells and T cells need to undergo an intricate process of differentiation. B cells differentiate into antibody-producing plasmablasts in germinal centers (GC), and upon pathogen clearance into long-lived memory B cells. Likewise, upon T cell priming, T cells differentiate into effector T cells, and upon pathogen clearance are maintained as memory T cells to ensure long-term production from recurring infections. In the past decennia, important insights have been obtained how B cells and T cell differentiate. In particular, the role of transcription factors and of metabolic regulators was extensively studied herein (11–14).

For appropriate lymphocyte differentiation, the regulators of differentiation processes must be produced at the right time and the right amount. In fact, gene dosage of transcription factors was shown to be key for B cell and T cell differentiation (15–18). This fine-tuning of gene expression is -at least in part -regulated by post-transcriptional events governed by RNA-binding proteins (RBPs) and non-coding RNAs (19,20). RBPs control a plethora of processes. They orchestrate RNA splicing, RNA polyadenylation and the subsequent export from the nucleus to the cytoplasm (21,22). RBPs can also modify the RNA (23). Furthermore, RBPs control mRNA localization, translation and stability. For instance, the RBPs ZFP36L1 and ZFP36L2 induce quiescence in developing B cells to allow for efficient B cell receptor rearrangement (24). ZFP36L1 is also required for the maintenance of the marginal-zone B cell compartment (25). For germinal center B cells that undergo cell cycle progression and affinity maturation, the expression of the RBP PTBP1 is key (26). In thymocytes, ZFP36L1 and ZFP36L2 dampen the DNA-damage response, which promotes their differentiation into mature T cells (27). In the periphery, Roquin suppresses T helper cell differentiation (28,29). Also m^6^A modifications are important for T helper cell differentiation, as evidenced in mice lacking the methyltransferase METTL3 in T cells (30).

Not only T cell differentiation, but also T cell effector function is tightly regulated by RBPs. Genetic ablation of the RBP Regnase*-*1 reprogrammed CD8^+^ T cells into long-lived effector CD8^+^ T cells, resulting in increased tumour control (31). In a patient, a nonsense-mutation in *REGNASE-1* resulted in hyperinflammation, including hypercytokinemia in T cells and monocytes (32). Another example is ZFP36L2, which blocks the cytokine production in memory CD8^+^ T cells from pre-formed mRNA in the absence of activation signals, thereby preventing aberrant production of effector molecules (33).

Even though these examples clearly highlight the importance of RBPs in regulating gene expression in lymphocytes, to date studies have addressed the contribution of individual RBPs. The overall expression profile of RBPs in primary human B cells and T cells is not well-documented, yet critical for our understanding of regulation of gene expression in lymphocytes.

In this study, we mapped the mRNA and protein expression of RBPs in primary human B cells and T cells. We observed clear differences of RBP expression levels between lymphocyte subsets. Furthermore, upon differentiation, the RBP expression profile significantly altered, which resulted in a shift of functional annotations of RBPs. Lastly, we identified an RBP signature that is specific for CD4^+^ T cells and CD8^+^ T cells with a high cytotoxic potential. In conclusion, RBP expression is lymphocyte-type specific and the RBP expression shows dynamic changes upon differentiation.

## RESULTS

### RNA-binding proteins are abundantly expressed in human B and T lymphocytes

To investigate the overall mRNA and protein expression of RBPs in human lymphocytes, we first generated a comprehensive list of RBPs. We included RBPs that were identified by RNA-interactome capture on multiple cell lines including, HEK293, HeLa-S3, MCF7, MCF10A, U2OS and Jurkat cells (34–36). This list was supplemented with computationally predicted RBPs based on the presence of a defined list of RNA-binding domains (RBDs) (36,37). This compiled list resulted in 3233 unique RBPs (Figure 1A, Supplementary Table 1).

**Figure 1:**
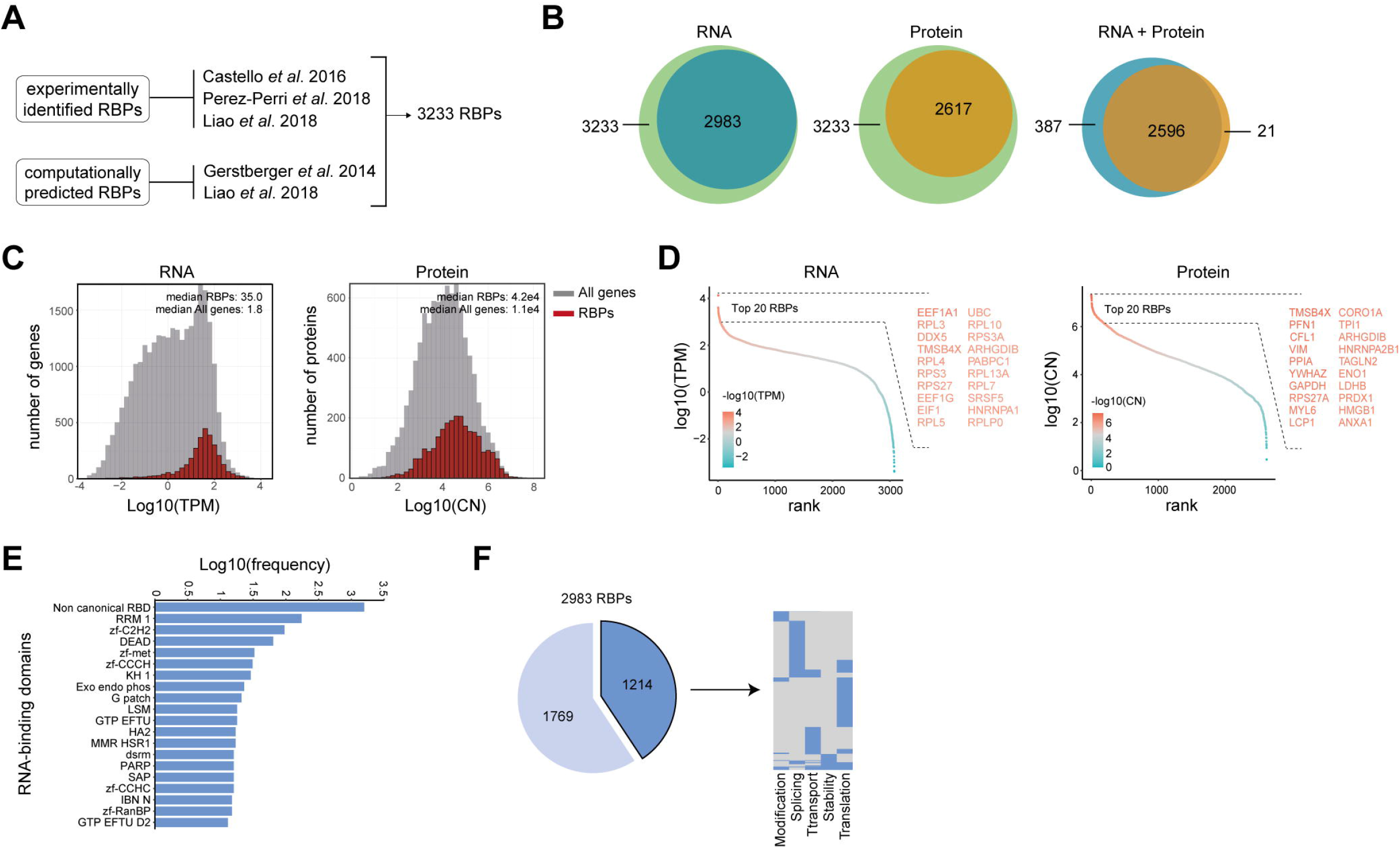
Characterization of RNA-binding protein expression in human lymphocytes. (**A**) Reference list of RNA binding proteins (RBPs) was generated by integrating experimentally validated RBPs (34–36) with computationally predicted RBPs based on the presence of a defined list of RNA-binding domains (36,37). (**B**) RBPs that are detected in human lymphocytes at RNA level (left panel), at protein level (middle panel), and at both RNA and protein level (right panel). RNA: n=3-4 donors. Protein: 4 donors. (>0.1 TPM) (**C**) RNA abundance in transcript per kilobase per million (TPM) and protein abundance in protein copy number (CN) for all genes (gray) and RBPs (red) in human B cells and T cells. (**D**) Expression levels of RBPs detected at RNA level (left panel) and at protein level (right panel) was ranked according to expression levels. Names of the top 20 expressed RBPs are indicated. (**E**) Frequency of RNA-binding domains among the 2983 RBPs that were detected at RNA level. (**F**) Left panel: RBPs detected at RNA level in human B and T lymphocytes that are annotated for RNA splicing, stability, subcellular localization of RNA, RNA modification, and translation (dark blue), or for other processes (light blue). Right panel: Distribution of RBPs annotated for the five RNA-related processes as indicated. Each line depicts one RBP. TPM: Transcripts per kilobase per million; CN: Protein copy number.

To define the global RBP gene expression in human B and T lymphocytes, we compiled previously published RNA-sequencing (RNA-seq) data on human CD19^+^ B cell, CD4^+^ T cell and CD8^+^ T cell subsets that were isolated from the blood of 3-4 healthy human donors (38). On average, 12.5×10^6^ reads per sample (range: 7.97×10^6^-19.15×10^6^ reads) could be mapped onto the human transcriptome. A total of 12,830 gene products (>0.1 TPM) were detected in all lymphocyte subsets combined. 2983 of the 3233 RBPs (92.3% of our reference list) were detected at the RNA level in human B and T lymphocytes (>0.1 TPM, Figure 1B), of which 2189 were identified in RNA-interactome capture studies and 794 were computationally predicted RBPs. The number of RBPs expressed at the RNA level in human B and T lymphocytes was similar to that of the epithelial cell line HeLa-S3 and the myelogenous leukemia cell line K562 cells, and overlapped for 90.1% (HeLa-S3: 2843 RBPs, K562: 2826 RBPs, Supplementary Figure 1A,B, (39)).

To calculate the number of RBPs expressed at the protein level in B and T lymphocytes, we used previously published mass spectrometry (MS) data of B cell and T cell subsets of 4 donors (40) that were similarly prepared and selected as the ones in the RNA-seq dataset we used (38). In total, 9436 proteins were identified in all B cell and T cell subsets combined, of which 96.8% (9136 proteins) were also expressed at the RNA level (Supplementary Figure 1C). Overall, 2617 RBPs (80.9% of our reference list) were detected at the protein level (Figure 1B), and 2596 RBPs (80.2%) were detected at both RNA and protein level (Figure 1B). This high overlap corroborated with the overall high expression levels of RBPs (Figure 1C; p-value: 6.2e-12 for RNA and 2.1e-12 for protein). The top 20 expressed RBPs at RNA level included several ribosomal proteins (*RPL3, RPL4, RPS3, RPS27*), translation-related proteins (*EEF1A1, EEF1G, EIF1*) and splicing-related proteins (*DDX5* (41), *SRSF5* (42), *HNRNPA1* (43); Figure 1D). At protein level, top 20 expressed RBPs included the RNA stability-related protein (VIM (44)), the splicing-related protein HNRNPA2B1 (45), the ribosomal protein RPS27A, and the moonlighting RBPs ENO1 and GAPDH (Figure 1D, (46,47)).

Next, we determined which RNA-binding domains (RBDs) are present in the detected RBPs. Using previously reported RBDs (Supplementary Table 1, (36,37)) and the protein families annotation database (Pfam; (48)), we detected a broad range of RBDs. (Supplementary Table 1). The top 20 RBDs included classical RBDs, such as the RNA-recognition motif (RRM; 5.8%, present in e.g. *CELF2, CNOT4, ELAVL2, HNRNPLL* and *PABPC1*) and the DEAD helicase motif (DEAD; 2.1%, present in e.g. *DDX1, DDX10, DHX16*). Also a variety of zinc-finger protein domains was found, including the zinc-finger C2H2 (zf C2H2; 3.2%, present in e.g. *ZNF638, ZMAT3*), zinc-finger metazoans (zf-met; 1.1%, present in e.g. *ZFR2, TUT1*), zinc-finger CCCH (zf-CCCH; 1.0%, present in e.g. *ZFP36L1, RBM27, ZC3H10*), zinc-finger CCHC (zf-CCHC; 0.5%, present in e.g. *ZCRB1, CPSF4*) and the zinc-finger Ran binding protein (zf-RanBP; 0.5%, present in e.g. *RBM5, RBM10*, Figure 1E). In addition, 52.8% of the RBPs contained non-canonical RNA-binding domains (1574 RBPs; Figure 1E), which were by and large present in experimentally identified RBPs (Supplementary Figure 1D). These included ribosomal proteins (*RPL18* and *RPL5*), the RNA processing molecule *DUSP11*, RNA splicing-related RBPs (*AHNAK, PCF11, SNIP1, SCAF11, SNRNP40*) and the exoribonuclease *EXOSC3*. A similar distribution of the top 20 RBDs was present in RBPs detected in HeLa-S3 and K562 cells (Supplementary Figure 1E), indicating that the RBD distribution is not a specific feature of lymphocytes.

RBPs regulate many processes, which includes RNA splicing, stability, subcellular localization of RNA, RNA modification, and translation (5,49). Using protein annotations from the human protein atlas database (50), we found that 1178 RBPs (41.7%) were annotated as regulators of at least one of these five RNA-related processes (Figure 1F, left panel), of which 24% were annotated for multiple RNA-related processes (Figure 1F, right panel). Combined, these data show that human lymphocytes express a wide variety of RBPs with a diverse set of RBDs.

### Human B cells and T cells have a distinct RBP signature

To determine whether and how RBP expression differed between B cells and T cells, we analyzed CD19^+^ B cells, CD4^+^ T cells, and CD8^+^ T cells separately in the RNA-seq and MS datasets employed in Figure 1. Overall, 2923 RBPs (97.0% of the total RBPs) and 2556 RBPs (97.6% of the total RBPs) were detected in all three subsets at the RNA and the protein level, respectively (Supplementary Figure 2A, B). Only few RBPs were identified in one cell type alone (Supplementary Figure 2A). In particular, B cells exclusively expressed members of the ribonuclease A super-family (*RNASE1, RNASE2 and RNASE3*) and the RBP *DAZL*. The RBPs *RBM24* and *PABPC3* were only detected in CD4^+^ T cells, and *NCBPL2* and *A1CF* were specifically expressed in CD8^+^ T cells (Supplementary Table 2). At the protein level, 36 RBPs were specifically detected in CD4^+^ T cells and CD8^+^ T cells (e.g. RC3H1, REXO1, HENMT1). KHDRBS2, RPL37, DHX32 were for instance exclusively detected in CD19^+^ B cells and CD4^+^ T cells and CPEB2 and TRMT44in CD4^+^ T cells alone (Supplementary Table 2).

**Figure 2:**
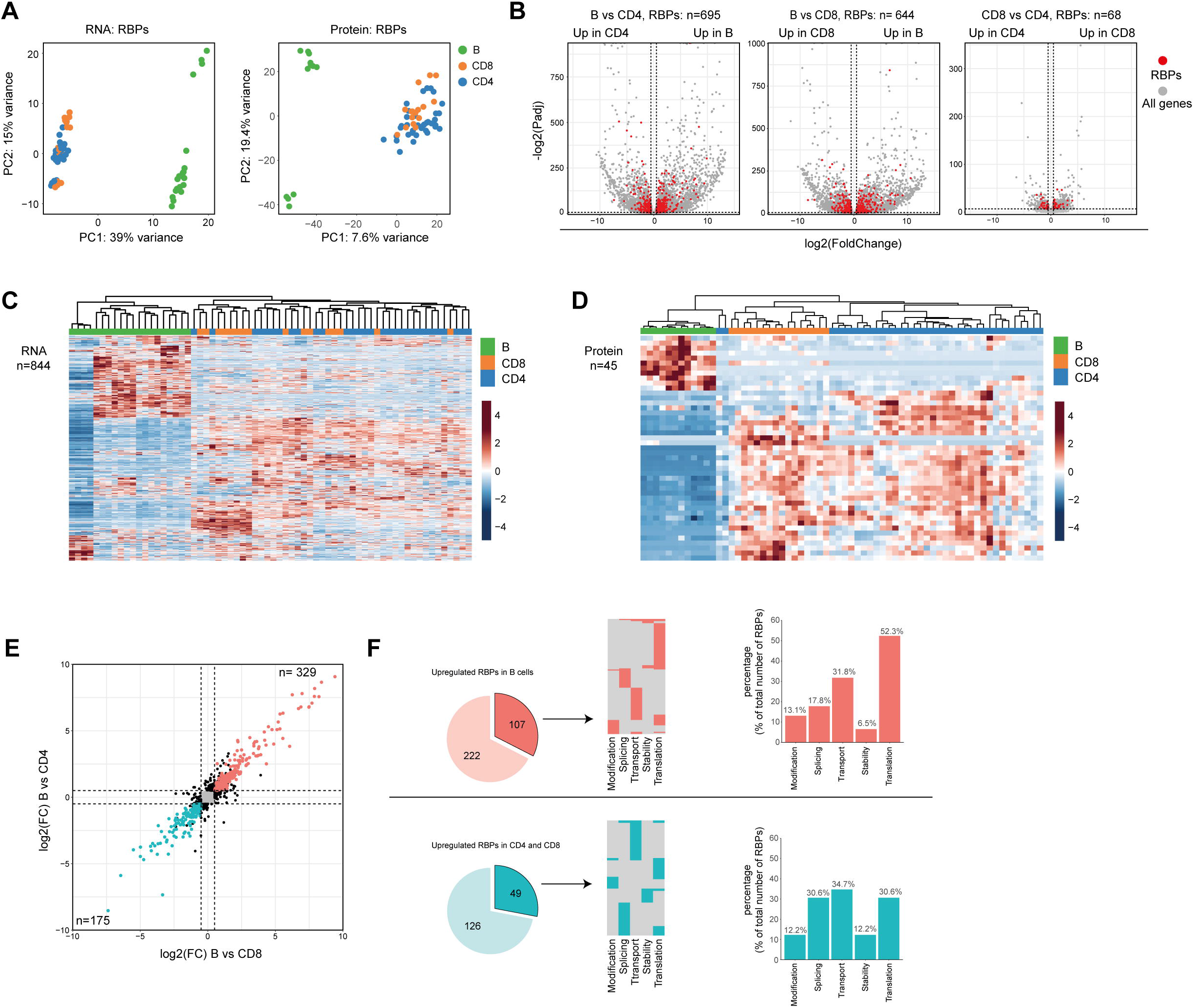
Differential RBP expression between human B cells and T cells. (**A**) Principal component analysis (PCA) of RBP RNA (left panel) and protein (right panel) expression in CD19^+^ B cells, CD4^+^ T cells and CD8^+^ T cells (n=4 donors). Each dot depicts one specific B cell or T cell subset from each donor. (**B**) Volcano plots of all differentially expressed genes (gray) and of differentially expressed RBPs (DE RBPs) (red) between CD19^+^ B cells, CD4^+^ T cells and CD8^+^ T cells (LFC>0.5, P-adjusted<0.01). (**C-D**) Heatmap of unsupervised clustering of DE RBPs at RNA (**C**) or at protein level (**D**) in CD19^+^ B cells, CD4^+^ T cells and CD8^+^ T cells. Each column corresponds to one T/B cell differentiation subset of a donor (n=4 donors). (**E**) Log2 Fold Change (LFC) of RBP mRNA expression between CD19^+^ B cells and CD4^+^ T cells (y-axis) and between CD19^+^ B cells and CD8^+^ T cells (x-axis). Red dots depict RBPs that are significantly upregulated in B cells, and blue dots indicate RBPs significantly upregulated in CD4^+^ T cells and CD8^+^ (significant in both comparisons, LFC>0.5, P-adjusted<0.01). (**F**) Left panels: RBPs annotated for RNA splicing, stability, subcellular localization of RNA, RNA modification, and translation (dark colors) or for other processes (light colors) that are upregulated in CD19^+^ B cells (top row) or T cells (bottom row) as defined in (**E**) Middle panels: relative distribution between the 5 specific RBP classes. Right panels: Percentage of RBPs annotated for the indicated RNA-related biological processes.

We next questioned whether the global RBP expression differed between the three lymphocyte subsets. Principal Component Analysis (PCA) revealed that the RBP mRNA and protein expression alone separates B cells from T cells just as effectively as a PCA performed on all genes (Figure 2A; Supplementary Figure 2C). Differential expression (DE) analysis on RBPs also revealed clear differences between B cells and T cells. 695 and 644 DE RBPs (82.3% and 76.3% of the DE RBPs) were found DE at the mRNA level between CD19^+^ B cells and CD4^+^ T cells, or CD8^+^ T cells, respectively (Figure 2B, Supplementary Table 2; LFC > 0.5; p-adjusted<0.01). CD4^+^ T cells and CD8^+^ T cells were more closely related, with only 68 DE RBPs (8.1% of all DE RBPs, Figure 2B, Supplementary Table 2). RBP expression at protein level showed similar trends, with 40 and 24 DE RBPs between CD19^+^ B cells and CD4^+^ T cells or CD8^+^ T cells, respectively, and only 6 DE RBPs between CD4^+^ T cells and CD8^+^ T cells (Supplementary Figure 2D, Supplementary Table 2; LFC > 0.5; p-adjusted<0.05). 84.4% of the DE RBPs at protein level were also DE RBPs at RNA level (Supplementary Figure 2E). Indeed, unsupervised clustering of the DE RBPs clearly distinguished B cell-from T cell-associated RBP clusters (Figure 2C, D).

To investigate the functional annotation of the DE RBPs, we focused on RBPs that were significantly higher expressed by either B cell or T cell populations (Figure 2E). We studied RBPs that are annotated regulators of RNA splicing, stability, subcellular localization of RNA, RNA modification, and translation, as defined by protein annotation from the human protein atlas database. This included 107 (32.5%) of the B cell-associated RBPs and 49 (28.0%) of the T cell-associated RBPs (Figure 2F, left panel). The majority of these RBPs were annotated for one function, and 17.8% and 16.3% for multiple functions for B cells and T cells, respectively (Figure 2F, middle panel). Interestingly, the relative distribution of RBPs annotated for these 5 RNA processes differed between B cells and T cells. Whereas 52.3% of RBPs in B cells were annotated for translation, this was only the case for 30.6% in T cells (Fig 2F, right panel). Conversely, only 17.8% was annotated for RNA splicing in B cells, but reached 30.6% in T cells (Fig 2F, right panel). In conclusion, the overt differential RBP expression between human B cells and T cells shown here possibly reflects a distinct distribution between different classes of RBPs.

### RBP expression changes upon B cell differentiation

Several B cell subsets can be found in the peripheral blood including naïve B cells, memory B cells and plasmablasts (Figure 3A). Whereas plasmablasts produce vast quantities of antibodies and are short-lived, memory B cells are long-lived and for the most part quiescent (51). We found that the phenotypical differences between these three B cell subsets is echoed in their RBP expression profile. We identified 1308 DE RBPs at RNA level, and 96 DE RBPs at protein level between naïve B cells, memory B cells and plasmablasts (Supplementary Figure 3A, B, Supplementary Table 3). In particular, although only 151 DE RBPs were found between naïve and memory B cells, plasmablasts showed a distinct RBP profile, with 1185 and 891 DE RBPs between naïve or memory B cells and plasmablasts, respectively (Supplementary Figure 3A, Supplementary Table 3). The top 20 DE RBPs at both the RNA and protein level spanned a wide range of abundance, and included *RRM2, APOBEC3B, METTL5* and *LGALS3* (Figure 3B, C). Hierarchical clustering of DE RBPs at RNA level revealed three clusters between B cell subsets (Figure 3D, Supplementary Table 3). 188 RBPs were highly expressed in plasmablasts (cluster 1), 541 RBPs were highly expressed in memory B cells (cluster 2) and cluster 3 with 579 RBPs were highly expressed in naïve and memory B cells, respectively (cluster 3). Hierarchical clustering on protein levels revealed similar differential RBP expression patterns (Figure 3C). Within these three clusters of DE RBPs at the RNA level, we isolated RBPs annotated for RNA splicing, stability, subcellular localization of RNA, RNA modification, and translation. This included 67 (35.6%) RBPs in cluster 1, 204 (44.4%) RBPs in cluster 2, and 214 (37%) RBPs in cluster 3 (Figure 3F, left panel). Again, 75-78% of the RBPs was annotated for one function (Figure 3F, middle panel), and the prime annotation of RBPs was translation in all three clusters (Figure 3F, right panel). Interestingly, whereas RBPs annotated for RNA splicing were also abundant in naïve and memory B cell subsets with 31.3% and 40.2%, respectively, plasmablasts (cluster 1) contained only 6.0% RBPs annotated for RNA splicing (Figure 3F, right panel). Instead, 44,8 % of RBPs expressed in plasmablasts annotated for RNA transport (Figure 3F, right panel). STRING-analysis on splicing-related RBPS from cluster 3 (naïve-memory B cells) revealed networks consisting of known splicing factors, such as the SR protein family members *SRSF1, SRSF4, SRSF3, SRSF6*, in addition to *NUDT21* and *HNRNPLL* (Figure 3G). For transport-annotated RBPs from cluster 1 (plasmablast), the interaction networks included the RBP *SLBP*, which regulates mRNA export (52), and the RBP *TST*, which regulates the transport of ribosomal RNA (Figure 3H, (53)). In conclusion, the RBP expression differs between B cell subsets, and involves different types of post-transcriptional regulatory functions.

**Figure 3:**
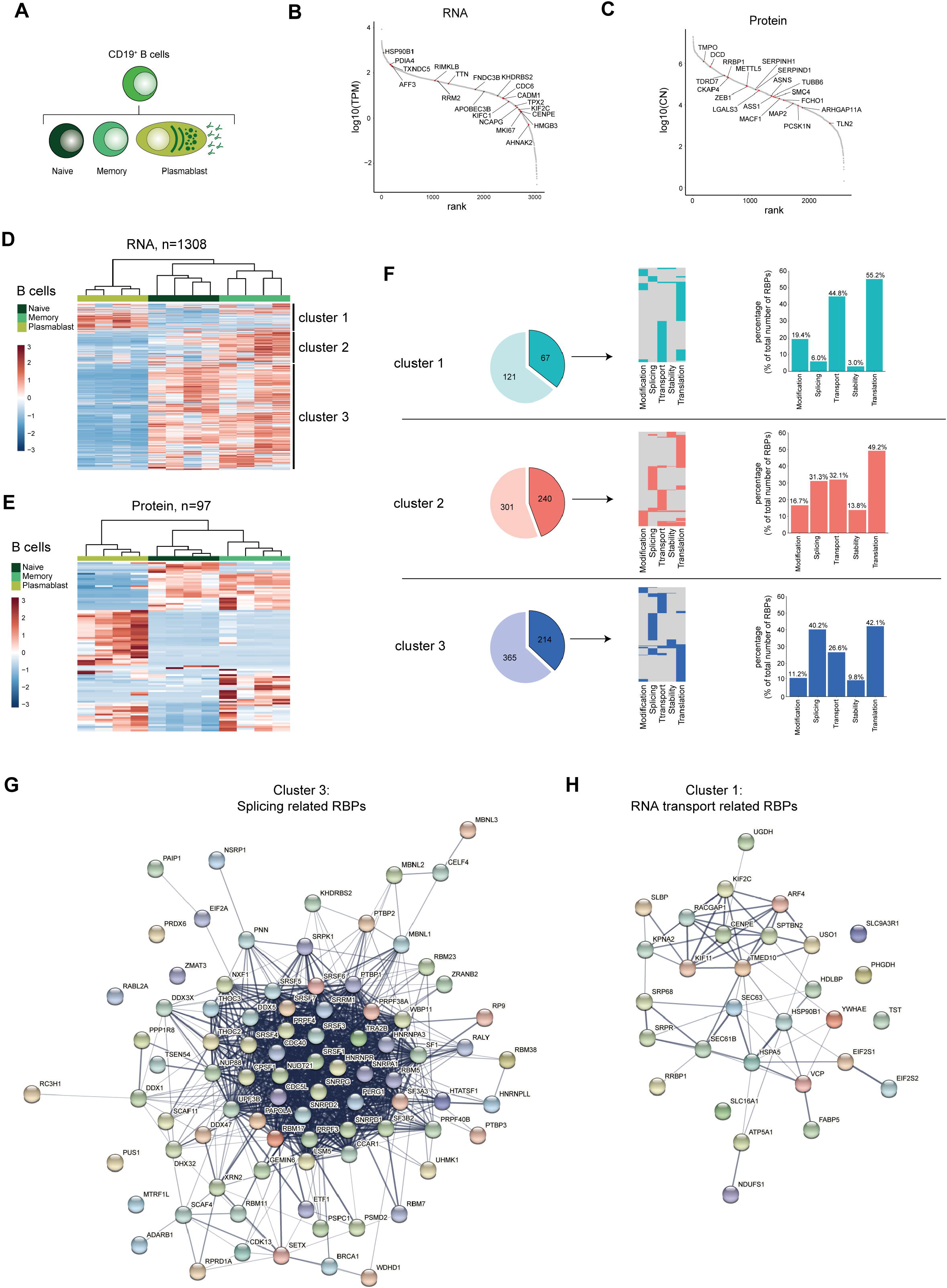
RBP expression alters upon B cell differentiation. (**A**) Diagram depicting the analysed CD19^+^ B cell subsets. (**B-C**) Expression levels of RBPs detected in B cells at RNA level (**B**) and at protein level (**C**), ranked according to expression levels. Red dots indicate the top 20 most differentially expressed RBPs, ranked on Log2 Fold Change). (**D-E**) Heatmap of unsupervised clustering of DE RBPs at mRNA (**D**) and at protein level (**E**) between naïve CD19^+^ B cells, memory CD19^+^ B cells and plasmablasts. n=4 donors. (**F**) Left panels: RBPs annotated for RNA splicing, stability, subcellular localization of RNA, RNA modification, and translation (dark colors) or for other processes (light colors) in the three clusters defined in (D). Middle panels: relative distribution between the 5 specific RBP classes. Right panels: Percentage of RBPs annotated for the indicated RNA-related biological processes. (**G-H**) String analysis on splicing-related RBPs (**G**) identified in cluster 3 and on RNA transport-associated RBPs (**H**) identified in cluster 1. TPM: Transcripts per kilobase per million; CN: Protein copy number.

### RBP expression changes upon CD4^+^ T cell differentiation

naïve T cells (Tnaive) undergo differentiation into effector T cells, which are rarely found in the peripheral blood of healthy donors (54). Rather, central memory (Tcm) and effector memory (Tem) CD4^+^ T cell subsets, which develop during the course of infections, are present in the blood and differentially contribute to recall responses upon recurring infections (Figure 4A; (55)). Similar to B cells, we find RBPs differentially expressed in the CD4^+^ T cell subsets Tnaive, Tcm and Tem at RNA (n=774), and at protein level (n=115; Supplementary Figure 4A, B; Supplementary Table 4). 48% of the DE RBPs at protein level are detected also at the RNA level (Supplementary Figure 4C). The top 20 DE RBPs included RBPs such as, *APOBEC3H* and *PAPBC3* (RNA level) and *OASL* and *ANXA2* (protein level, Figure 4B, C).

**Figure 4:**
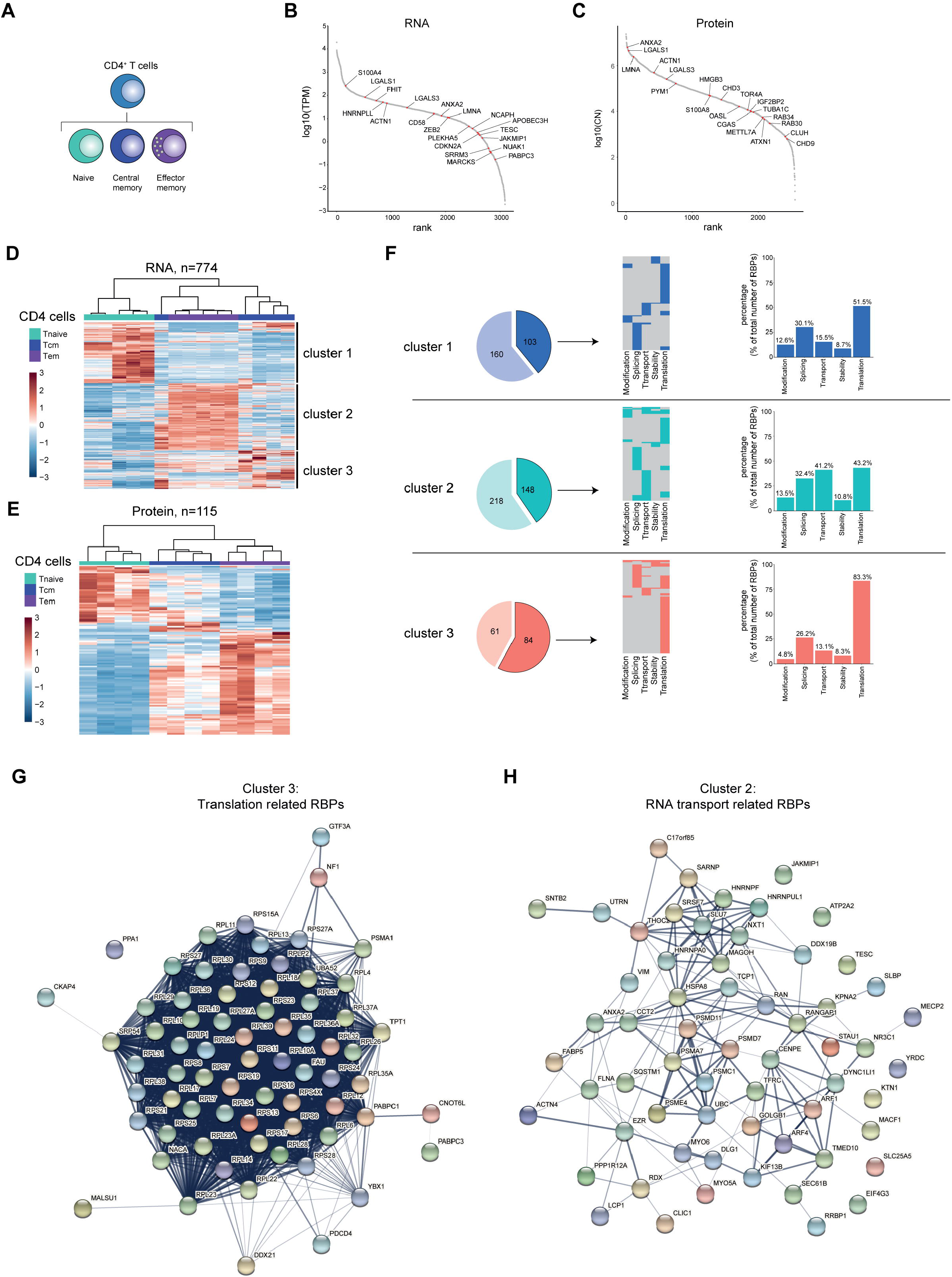
RBP expression alters upon CD4^+^ T cell differentiation. (**A**) Diagram depicting the analysed CD4^+^ T cell subsets. (**B-C**) Expression levels of RBPs detected at RNA (**B**) or protein level (**C**) in human CD4^+^ T cells ranked according to expression levels. Red dots indicate top 20 most differentially expressed RBPs based on Log2 Fold Change. (**D-E**) Unsupervised clustering of DE RBPs at RNA (**D**) or at protein level (**E**) between naïve (Tnaive), central memory (Tcm) and effector memory (Tem) CD4^+^ T cells depicted in a heatmap. RNA: n=5 donors, protein: n=4 donors. (**F**) Left panels: RBPs annotated for RNA splicing, stability, subcellular localization of RNA, RNA modification, and translation (dark colors) or for other processes (light colors) in the three clusters defined in (D). Middle panels: relative distribution between the 5 indicated RBP classes. Right panels: Percentage of RBPs annotated for indicated RNA-related biological processes. (**G-H**) String analysis on translation-related RBPs (**G**) identified in cluster 3 and on RNA transport-associated RBPs (**H**) identified in cluster 2 . TPM: Transcripts per kilobase per million; CN: Protein copy number.

Hierarchical clustering of the DE RBPs revealed three clusters (Figure 4D, E), with cluster 1 (Tnaive) containing 145 RBPs, cluster 2 (Tem) containing 366 RBPs, and cluster 3 (Tcm) containing 263 RBPs (Supplementary Table 4). This differential expression of RBPs was also apparent at the protein level (Figure 4E, 115 DE RBPs, Supplementary Table 4). To gain more insights into the biological processes of the RBPs in the different clusters, we performed Gene Ontology (GO) analysis. Whereas metabolic processes were enriched in all three clusters, cluster 2 and 3 were enriched for GO-terms associated with translation (translation initiation, cytoplasmic translation) and with RNA transport (Supplementary Figure 4D). Cluster 2 also showed a moderate enrichment for RBPs associated with regulation of RNA stability (RNA destabilization, 3’-UTR-mediated mRNA destabilization; Supplementary Figure 4D).

When we specifically isolated RBPs annotated for RNA splicing, stability, subcellular localization of RNA, RNA modification, and translation, we found that 103 (39.2%) RBPs of the DE RBPs belong to these 5 RBP classes in cluster 1, 148 RBPs in cluster 2 (40.4%) and 84 RBPs in cluster 3 (57.9%) (Figure 4F, left panels). Only a fraction of RBPs is associated with more than one of these functions (cluster 1: 17.5%, cluster 2: 26.4%, cluster 3: 25%; Figure 4F, middle panel). When CD4^+^ T cells differentiate, the relative distribution of functional RBP annotation alters. 83.3% of the RBPs associated with the 5 RBP classes were linked to translation in cluster 3 (Tcm), compared to 51.5% and 43.2% in cluster 1 (Tnaive) and cluster 2 (Tem), respectively (Figure 4F, right panels). Conversely, in cluster 2 (Tem), the percentage of RBPs annotated for RNA transport are with 41.2% primarily found in cluster 2 and much less so in cluster 2 and 3 with 15.5% and 13.1%, respectively (Figure 4F, right panels). STRING-analysis on the translation-related RBPs of cluster 3 revealed an enrichment of 53 ribosomal proteins and of other translation-associated RBPs, such as *PABPC1, YBX1* and *FAU* (Figure 4G). The RNA-transport-related RBPs of cluster 2 included the mRNA export-associated RBPs *DDX19B, SARNP, MAGOH* and *THOC2* (Figure 4H). Combined, our findings reveal that the RBP expression landscape changes throughout CD4^+^ T cell differentiation, which results in a relative enrichment of specific RBP classes in different CD4^+^ T cell subsets.

### RBP expression changes upon CD8^+^ T cell differentiation

We then focused on the CD8^+^ T cell differentiation subsets. Our dataset also included effector CD8^+^ T cells (Teff), which was included in the analysis, in addition to Tnaive, Tcm and Tem CD8^+^ T cell subsets (Figure 5A). 707 RBPs were differentially expressed at the RNA level between Tnaive, Tcm and Tem and Teff, and 115 RBPs at the protein level (Supplementary Figure 5A, B; Supplementary Table 5). 44.1% of DE RBPs at protein level were also detected at the RNA level (Supplementary Figure 5C). The top 20 DE RBPs included RBPs like *JAKMIP1* and *OASL* (RNA level) and EIF4EBP3 and FLNB (protein level, Figure 5B, C). Hierarchical clustering of RBP expression resulted 3 clusters (Figure 5D). Cluster 1 contained 297 RBPs highly expressed in Tem and Teff CD8^+^ T cells (Figure 5B, Supplementary Table 5). Cluster 2 (176 RBPs) also included Tem an Teff CD8^+^ cells, and to a lesser extent in Tcm cells. Cluster 3 (234 RBPs) included primarily Tnaive cells, but also Tcm cells (Figure 5B, Supplementary Table 5). Similar clusters were identified at RBP protein level (Figure 5E, 177 DE RBPs, Supplementary Table 5).

**Figure 5:**
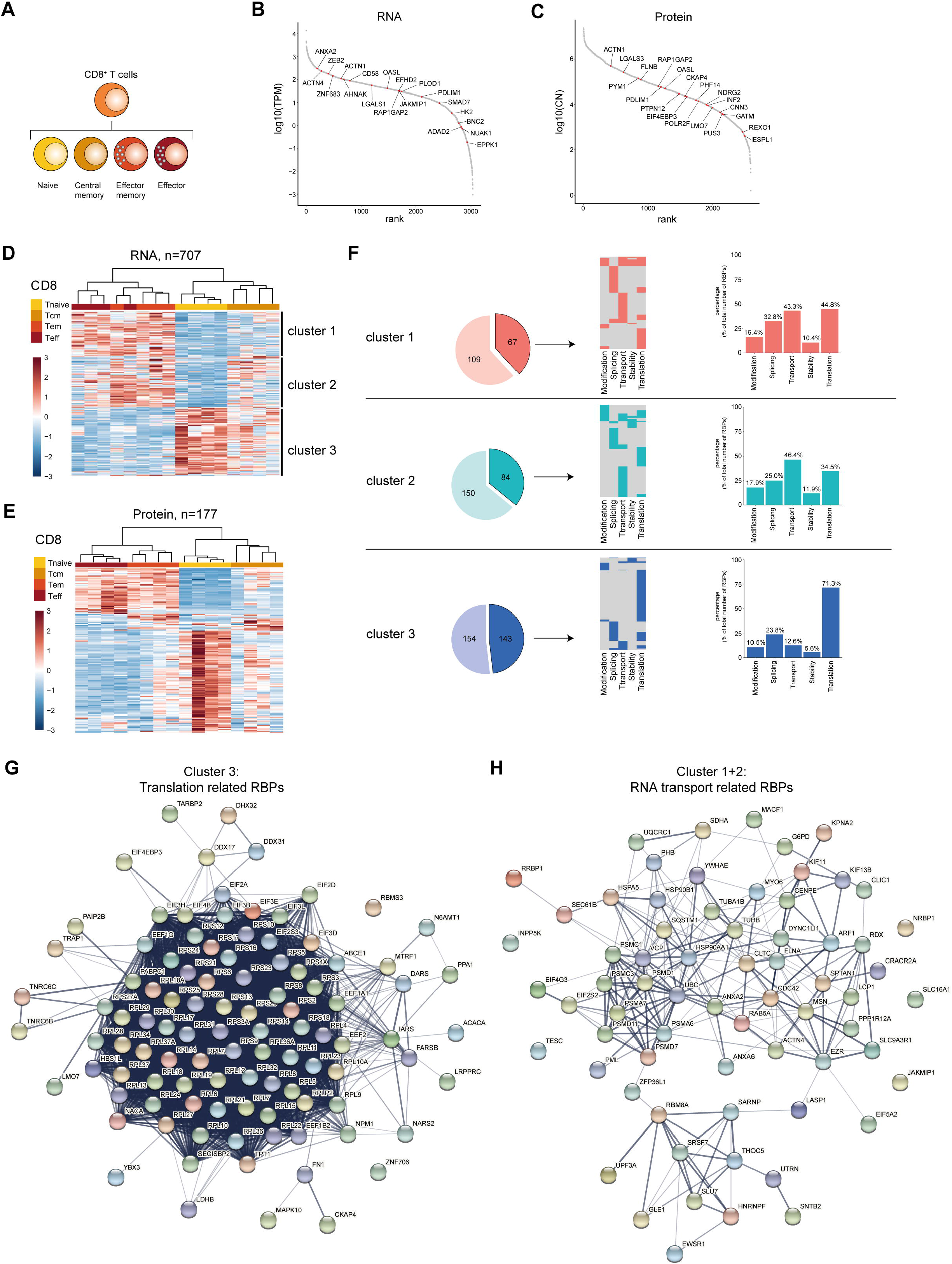
RBP expression alters upon CD8^+^ T cell differentiation. (**A**) Diagram depicting the analysed CD8^+^ T cell subsets. (**B-C**) Expression levels of RBPs detected at RNA (**B**) and at protein level (**C**) in human CD8^+^ T cells ranked according to expression levels. Red dots indicate top 20 most differentially expressed RBPs based on Log2 Fold Change. (**D-E**) Unsupervised clustering of DE RBPs at RNA (**D**) or protein level (**E**) between naïve (Tnaive), central memory (Tcm), effector memory (Tem) and effector (Teff) CD8^+^ T cells depicted in a heatmap. n=4 donors. (**F**) Left panels: RBPs annotated for RNA splicing, stability, subcellular localization of RNA, RNA modification, and translation (dark colors) or for other processes (light colors) in the three clusters defined in (D). Middle panels: relative distribution between the 5 indicated RBP classes. Right panels: Percentage of RBPs annotated for indicated RNA-related biological processes. (**G-H**) String analysis on translation-related RBPs (**G**) identified in cluster 3 and on RNA transport-associated RBPs (**H**) identified in clusters 2 and 3. TPM: Transcripts per kilobase per million; CN: Protein copy number.

In CD8^+^ T cell subsets, 67 RBPs in cluster 1 (38.1%), 84 RBPs in cluster 2 (35.9%), and 143 RBPs in cluster 3 (48.1%) were annotated as regulators of RNA splicing, stability, subcellular localization of RNA, RNA modification, or translation (Figure 5F, left panels), with a minority of RBPs (20-28%) linked to multiple functions (Figure 5F, middle panels). We found that cluster 3 was relatively enriched for translation-associated RBPs (71.3%), compared to cluster 1 and cluster 2 with 34.5% and 44.8%, respectively (Figure 5F, right panels). Conversely, cluster 1 and 2 were enriched for RBPs associated with RNA transport (cluster 1: 43.3%, cluster 2: 46.4%), and this RBP class was 12.6% only minor in cluster 3 (Figure 5F, right panels). STRING-analysis on translation-associated RBPs from cluster 3 revealed the interaction network between 56 ribosomal proteins and 10 eukaryotic translation initiation factors (Figure 5G). The RNA-transport associated RBPs in cluster 2 and 3 included RBPs involved in RNA export (*THOC5* (56), *RBM8A* (57), *SARNP* (58), Figure 5H).

Gene Ontology (GO) analysis on the DE RBPs also showed in cluster 3 -in addition to catabolic processes -an enrichment of GO-terms associated with translation, i.e. cytoplasmic translation, translation initiation and positive regulation of translation (Supplementary Figure 5D). Cluster 1 displayed a moderate enrichment for GO-terms related to RNA stability (3’-UTR-mediated mRNA destabilization, regulation of mRNA stability, Supplementary Figure 5D). In conclusion, CD8^+^ T cells change their RBP expression landscape throughout differentiation, with specific RBP classes enriched in different CD8^+^ T cell subsets.

### Specific RBP expression associates with T cell cytotoxicity

T cells can acquire cytotoxic function when they differentiate into effector cells. Importantly, whereas CD8^+^ T cells are generally classified as cytotoxic, not all CD8^+^ T cells display cytotoxic features (59–61). Conversely, a subset of human CD4^+^ T cells also shows cytolytic features (62–64). We therefore sought to identify RBPs that were associated with a high cytotoxic capacity in human CD8^+^ T cells and CD4^+^ T cells. As source of T cells, we used previously published single-cell RNA-seq (scRNA-seq) data on blood-derived human CD8^+^ and CD4^+^ T cells (65–67).

Because only memory and effector T cells can be cytotoxic, we excluded naïve T cells from our analysis based on their high gene expression of *CCR7, LEF1* and *SELL* (Supplementary Figure 6A-D). We then identified and integrated the expression of 8 cytotoxic genes (9) i.e. *FGFBP2, GZMB, GZMH, PRF1, NKG7, CX3CR1, GNLY* and *ADGRG1* into a *cytotoxic score* (*see methods*; Figure 6A, Supplementary Figure 6E, F). Dimensional reduction analysis revealed that CD8^+^ and CD4^+^ T cells with a low (bottom 10%) or high cytotoxic score (top 10%) form two distinct clusters (Figure 6B, C). High expression of *ITGB1* in CD8^+^ and CD4^+^ T cells with a high cytotoxic score confirmed the selection for cytotoxic T cells ((59,62), Figure 6D, E). In addition, 16 RBPs were significantly upregulated in CD8^+^ T cells with a low cytotoxic score, whereas 36 RBPs were preferentially expressed in CD8^+^ T cells with a high cytotoxic score (Figure 6D, Supplementary Table 6; LFC>0.5; P-adjusted<0.01). Likewise, 87 RBPs and 41 RBPs were upregulated in CD4^+^ T cells with a low and with a high cytotoxic score, respectively (Figure 6E, Supplementary Table 6).

**Figure 6:**
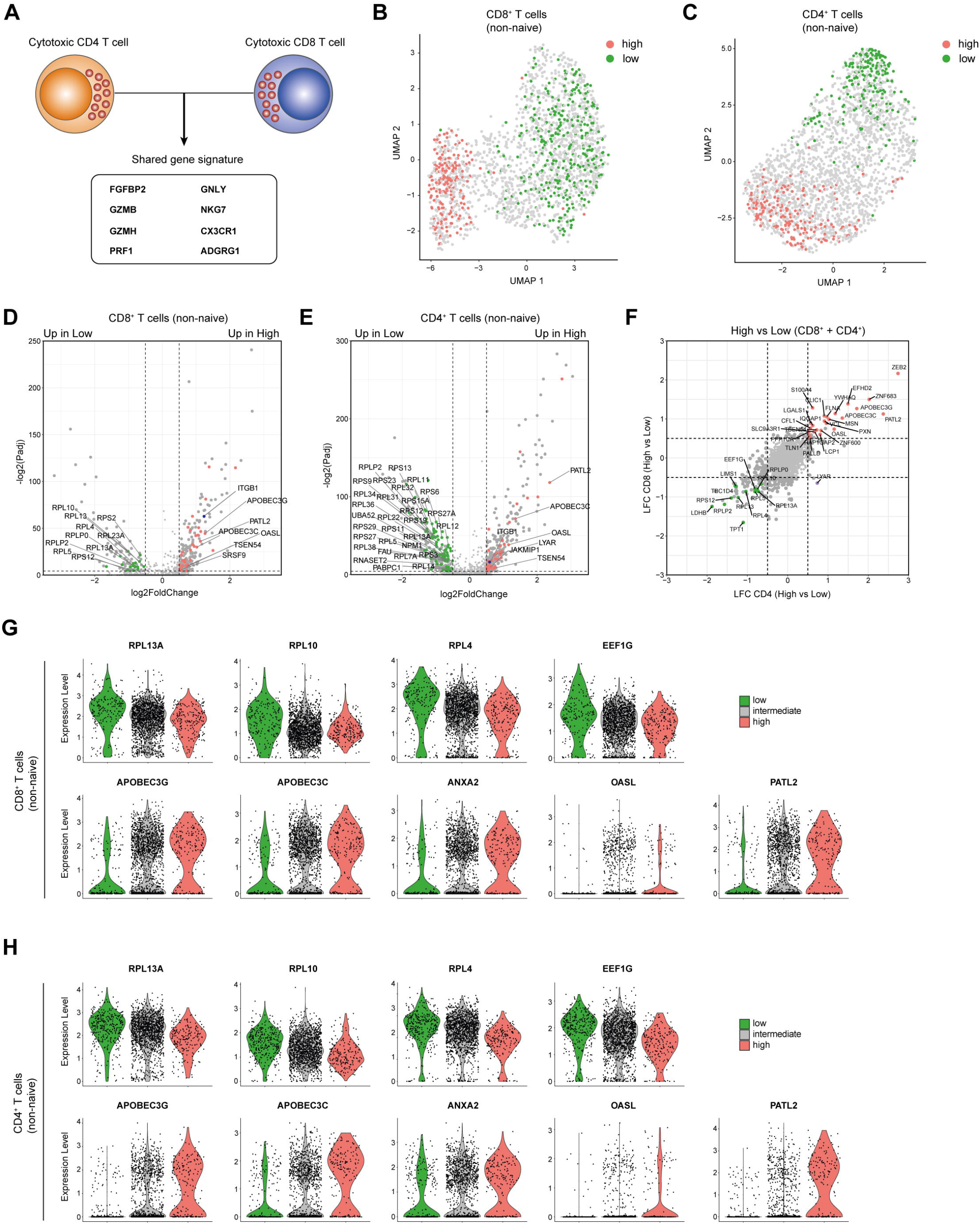
Cytotoxic CD4^+^ and CD8^+^ T cells share the RBP expression profile. (**A**) Diagram indicating the cytotoxic gene signature shared by CD4^+^ T cells and CD8^+^ T cells as defined in Supplementary Figure 6E,F. (**B, C**) Uniform Manifold Approximation and Projection (UMAP) plot on non-naïve CD8^+^ T cells (B) and CD4^+^ T cells (C) with a high (top 10%, red) or low (bottom 10%, green) cytotoxic score. (**D, E**) Volcano plot d DE RBPs (red) and other genes (gray) between non-naïve CD8^+^ T cells (D) and CD4^+^ T cells (E). Blue dot depicts *ITGB1*. (**F**) Log2 Fold Change values for RBPs with a high or low cytotoxic score of non-naïve CD4^+^ T cells (y-axis) and non-naïve CD8^+^ T cells (x-axis). Red and green dots indicate DE RBPs associated with a high and low cytotoxic score in both T cell types (LFC>0.5 P-adjusted<0.05). (**G, H**) Violin plots depicting expression levels and expression density of selected RBPs in CD8^+^ T cells (G) or CD4^+^ T cells (H) with a high (top 10%), intermediate (10-90%) or low (bottom 10%) cytotoxic score.

Intriguingly, the differential RBP expression between CD8^+^ and CD4^+^ T cells with a low or high cytotoxic score was strikingly similar (Figure 6F). 13 RBPs were upregulated in CD8^+^ and CD4^+^ T cells with a low cytotoxic score (Figure 6F), which included 8 ribosomal proteins, e.g. *RPL13A, RPL10* and *RPL4* which are accessory to the translation regulation (68), and the translation initiation factor *EEF1G* (Figure 6G, H). 25 RBPs that were upregulated in both CD8^+^ and CD4^+^ T cells with a high cytotoxic score (Figure 6F) included cytidine deaminases APOBEC3G and APOBEC3C, the poly(G) binding protein ANXA2 (69), the viral dsRNA binder OASL (70) and the translational repressor PATL2 (Figure 6G, H, (71)). In summary, CD8^+^ and CD4^+^ T cells with a high cytotoxic potential express a specific set of RBPs.

## Discussion

In this report, we provide a comprehensive RBP expression map based on the transcriptome and proteome of human B cells and T cells. We found that the overall expression pattern of RBPs is remarkably well conserved. It overlaps for >90% with that of HeLa-S3 cells and K562 cells. Nonetheless, differential expression analysis of RBPs clearly distinguishes B cells from T cells. RBPs also alter their expression profile upon differentiation. This finding suggests that -similar to what is observed for transcription factors (15,16) -the relative abundance of RBPs defines the fate of RNA and of translation into proteins, and thus the differentiation status of lymphocyte subsets.

Intriguingly, the differential expression of RBPs upon lymphocyte differentiation resulted in a shift of functional annotations of the expressed RBPs. For instance, plasmablasts are enriched for RBPs annotated for RNA transport, a feature that may support their antibody producing function. Effector and effector memory type CD4^+^ and CD8^+^ T cells also showed a preference of RBPs annotated for RNA transport, albeit to a lesser extent. Conversely, central memory CD4^+^ T cells, and naïve and central memory CD8^+^ T cells preferentially express RBPs that are annotated for translation regulation. This finding is in line with recent studies that indicated a tight gene-specific regulation of translation in naïve T cells (72–75), and the concept of translational preparedness of naïve and memory T cells (72). It is therefore tempting to speculate that the enrichment for RBPs involved in translation regulation we find here contributes to the translation control in naïve and memory T cells.

Also, cytotoxic T cells display a unique RBP expression profile. Compared to non-cytotoxic T cells, cytotoxic T cells express lower mRNA levels of several ribosomal proteins. Whether the differences at the mRNA level for ribosomal proteins is also reflected at the protein level is still unknown. In addition, whether some of these ribosomal proteins display any transcript specificity, as was shown for RPL10A and RPS25 (68), remains to be defined. In addition, we noticed an increased expression of the mRNA cytidine deaminases APOBEC3C and APOBEC3G. Previous studies reported that APOBEC expression increases upon T cell activation (76), which was primarily associated with viral restriction (77). It is also conceivable that increased APOBEC expression is involved in regulating the fate of endogenous mRNAs in cytotoxic T cells. Interestingly, the specific RBP profile linked to cytotoxicity is shared by CD4^+^ T cells and CD8^+^ T cells, a feature which may point to a similar differentiation program towards cytotoxicity.

RNA-binding proteins are critical mediators in shaping lymphocyte differentiation and effector function (24–28,30–33). The RBP expression map we provide here could help to further dissect the role of RBPs in B cell and T cell differentiation and function. It is important to note that the RBP expression map we provide here primarily serves as a resource, and thus as a starting point for uncovering the RBP-mediated regulation in lymphocyte differentiation. RBP expression by itself cannot be interpreted as direct interaction of RBPs with RNA. In fact, RBP interactions with RNAs can be highly versatile and are subject to rapid changes upon extrinsic signals. For instance, the RBP ZFP36L2 is expressed to a similar extent in memory T cells and re-activated T cells, yet only blocks translation in resting memory T cells (33). Similarly, a large fraction of ribosomal proteins does not interact with ribosomal RNA. The mode of action of these non-ribosomal RNA binding RPs is to date enigmatic and requires further investigation. Lastly, in addition to classical RBPs, recent studies have revealed the presence of enigmatic RBPs, which are primarily annotated for other cellular functions. This is exemplified by metabolic enzymes from the TCA cycle (78,79). Their relative contribution to RNA regulation during lymphocyte differentiation and effector function is yet to be experimentally confirmed. Nonetheless, the role of RBPs in genetic diseases is becoming appreciated (80), and defining the RBP expression presented in the study presented here may contribute to deciphering dysregulated RBP expression and function also in immune-related diseases.

## Material and Methods

### Data sets

Raw RNA-sequencing (RNA-seq) data were retrieved from the gene expression omnibus repository (GEO, NCBI) or from the European Nucleotide Archive (ENA). Data from CD19^+^ B cells and from B cell differentiation subsets (n=4 donors with 4 B cell populations each), from CD4^+^ T cells (n=3-4 donors with 7-8 CD4^+^ T cell populations each) and CD8^+^ T cells and respective differentiation subsets (n=4 donors with 4 CD8^+^ T cell populations each) were retrieved from Monaco *et al*. ((38); accession number: GSE107011)). RNA seq data to of CD4^+^ T cell subsets (n=5 donors with 3 CD4^+^ T cell populations each) were retrieved from Ranzani *et al*. ((81); PRJEB5468). RNA-seq data of HeLa-S3 cells and K562 cells were obtained from Martinez *et al*. ((39); GSE125218). Quantitative mass spectrometry (MS) data of CD19^+^ B cells, CD4^+^ and CD8^+^ T cells and the respective differentiation subsets were retrieved from Rieckmann *et al*. ((40) n=4 donors). Lymphocyte subsets in the RNA-seq and MS data sets were prepared as follows: CD19^+^ B cells: naïve (RNA-seq: CD27^-^ IgD^+^, MS: CD27^-^ Mitotracker^-^), memory (RNA-seq: CD27^+^ CD38^-^ IgD^-^, MS: CD27^+^ CD38^-^ Mitotracker^-^) and plasmablasts (RNA-seq: CD27^+^ CD38^+^ IgD^-^, MS: CD27^+^ CD38^+^ Mitotracker^-^); CD4^+^ T cells: naïve (RNA-seq: CCR7^+^ CD45RA^+^ CD45RO^-^, MS: CCR7^+^ CD45RA^+^), memory (RNA-seq: CCR7^+^ CD45RA^-^ CD45RO^+^, MS: CCR7^+^ CD45RA^-^), and effector memory (RNA-seq: CCR7^-^ CD45RA^-^ CD45RO^+^, MS: CCR7^+^ CD45RA^-^); CD8^+^ T cells: naïve (CCR7^+^ CD45RA^+^), memory (CCR7^+^ CD45RA^-^), effector memory (CCR7^-^ CD45RA^-^) and effector CD8^+^ T cells (CCR7^-^ CD45RA^+^).

Single-cell RNA-sequencing (scRNA-seq) data of blood-derived CD4^+^ and CD8^+^ T cells were retrieved from the GEO repository: Zheng *et al*. (67); GSE98638, Guo *et al*. (65); GSE99254, Zhang *et al*. (82); GSE108989.

### RBP reference list

The list of annotated human RBPs was created by aggregating published data of RNA interaction capture assays that were performed on HEK293, HeLa, MCF7, MCF10A, U2OS and Jurkat cells (34–36), which resulted in a list of 2355 RBPs. This list was supplemented with 975 computationally identified RBPs from Gerstberger *et al*. (37), and the EuRBP-DB ((36), http://eurbpdb.syshospital.org/, accessed on 19-11-2019. This RBP list with 3333 proteins was manually curated to exclude histones (18 histones), possible contaminants (ITGA1 and ITGB1), and mitochondrial RBPs (80 RBPs), resulting in a list of 3233 RBPs.

### RNA-sequencing analysis

RNA-sequencing reads were quasi-mapped using Salmon (version 1.0, (83)) onto the human coding transcriptome GRCh38 from Gencode (v36, May 2020). Of the CD4^+^ T cell, CD8^+^ T cell and CD19^+^ B cell samples retrieved from Monaco *et al*. (38), an average of 12.5 × 10^6^ reads was quasi-mapped onto the human coding transcriptome. For the CD4^+^ T cell differentiation samples, retrieved from Ranzani *et al*. (81), an average of 11.4 ⨯ 10^6^ reads was quasi-mapped onto the human coding transcriptome. Transcript-level estimates were imported and summarized to the gene-level by using the tximport function (tximport package, version 1.16.1 (84)). To define the overall expression of RBPs subsets were grouped together as indicated. Differential gene expression analysis was performed using DESeq2 (version 1.28.1 (85)). Genes were considered differentially expressed with an absolute log2 fold change (LFC) >0.5 and a *p*-adjusted <0.01. For differential gene expression analysis of total B cells, CD4^+^, and CD8^+^ T cells populations, we averaged the RNA-seq counts of differentiation subsets per donor. TPM (transcript per kilobase per million) counts were calculated by Salmon and used for plotting. Of note, TPM counts are corrected for library depth, library size, and transcript length, and thereby allow a fair comparison between populations. The number of detected RBPs per cell type, and the RBP expression rank were based on averaged TPM per cell type. Top 20 differentially expressed proteins were identified based on the log2 fold change values.

### Single cell RNA-seq data analysis

ScRNA-seq datasets were analysed using Seurat (version 4.0.1;(86)). Count matrices of (65,67,82) were filtered for “PTC”, corresponding to peripheral blood-derived CD8^+^ T cells (CD3^+^CD8^+^). To identify conventional blood-derived CD4^+^ T cells, count matrices of references (65,67,82) were filtered for “PTH” (CD3^+^CD4^+^CD25^-^) and “PTY” (CD3^+^CD4^+^CD25^int^). To correct for dataset specific effects from the three individual scRNA-seq datasets, we employed a published scRNA-seq data integration method (87). The inter-individual donor batch-effect was corrected using the *vars*.*to*.*regress* argument in SCTransform (Seurat v4). Unsupervised clustering was performed on Uniform Manifold Approximation and Project (UMAP) dimensional reduction using the top 30 principal components (PCs). Cells expressing high levels of naive T cell associated genes like *CCR7, LEF1* and *SELL* (88,89) were excluded from downstream analysis. Differential gene expression analysis was performed using the Model-based Analysis of Single-cell Transcriptomics (MAST) test (90). Genes were considered differentially expressed based on a *p*-adjusted < 0.05 and an absolute log2 fold change > 0.5.

### Cytotoxic score calculation

The cytotoxic score of CD4^+^ T cells and CD8^+^ T cells was obtained from the scRNA-seq data by selecting for the top 7 most correlated (Pearson’s correlation) genes with *FGFBP2* expression (9). (Supplementary Figure 6A, B). To obtain the cytotoxic score for each cell, a Z-score of expression for each of the 8 cytotoxic genes (*FGFBP2, GZMB, GZMH, PRF1, NKG7, CX3CR1, GNLY* and *ADGRG1)* was calculated for the whole dataset. Z-scores from all 8 genes were averaged per cell and served as the cytotoxic score. Cells with high (top 10%), intermediate (10-90%) or low (bottom 10%) cytotoxic score were selected and used for analysis.

### Mass spectrometry analysis

Differential protein expression analysis was performed with Differential Enrichment analysis of Proteomics data (DEP) (version 1.12.0, (91)) on label-free quantification (LFQ) values. Protein copy numbers (CN) were filtered for expression levels (CN > 1). The number of RBPs detected among the different cell types was based on averaged CN values across cell types obtained from 4 donors. RBP rankings according to protein abundance was performed by using averaged CN values per cell type. The top 20 differentially expressed proteins were identified based on the log2 fold change values.

### RBD annotation

RNA-binding domain names were obtained from Gerstberger *et al*. (2014) and Liao *et al*. (2020). The existence of each RNA-binding domain was verified and updated based on information present in the protein families database (Pfam, (48)) (Supplementary Table 1). Proteins containing RNA-binding domains were obtained from the PFAM database. When RBPs contained more than one RBD, each RBD was counted and included in the analysis. Human Protein Atlas annotations (HPA, https://www.proteinatlas.org on 28-03-2021; (50)) were used to classify proteins associated with RNA modification (keywords: “RNA AND Modification”), RNA splicing (keywords: “Spliceosome”), RNA stability (keywords: “RNA AND Stability”), RNA transport (keywords: “RNA AND Transport”), and Translation (keywords: “Translation”). Protein-protein association networks were generated using the STRING database (https://string-db.org/, (92)). Gene ontology analysis was performed with the Panther database (version 16.0, (93)) on differentially expressed RBPs. A statistical overrepresentation test (Fisher’s exact with FDR multiple test correction) was performed with a reference list composed of all Homo Sapiens genes in the database. Overrepresented GO terms (FDR<0.05) were filtered for RNA-related functions. Full lists of overrepresented GO terms are provided in Supplementary Table 7.

### Plots and graphs

Plots and graphs were generated using ggplot2 (version 3.3; (94)). Principal components analysis was performed using the plotPCA function from DESeq2 (85). Heatmaps were generated using the Pheatmap package (version 1.0.12; (95)) in R (version 4.0.3). Venn diagrams were generated using the Venn diagram tool from the University of Gent (accessed at http://bioinformatics.psb.ugent.be/webtools/Venn/).

## Acknowledgements

The authors would like to thank Drs. B. Popovic and I. Foskolou for critical reading of the manuscript.

---

**Supplementary Figure 1.**
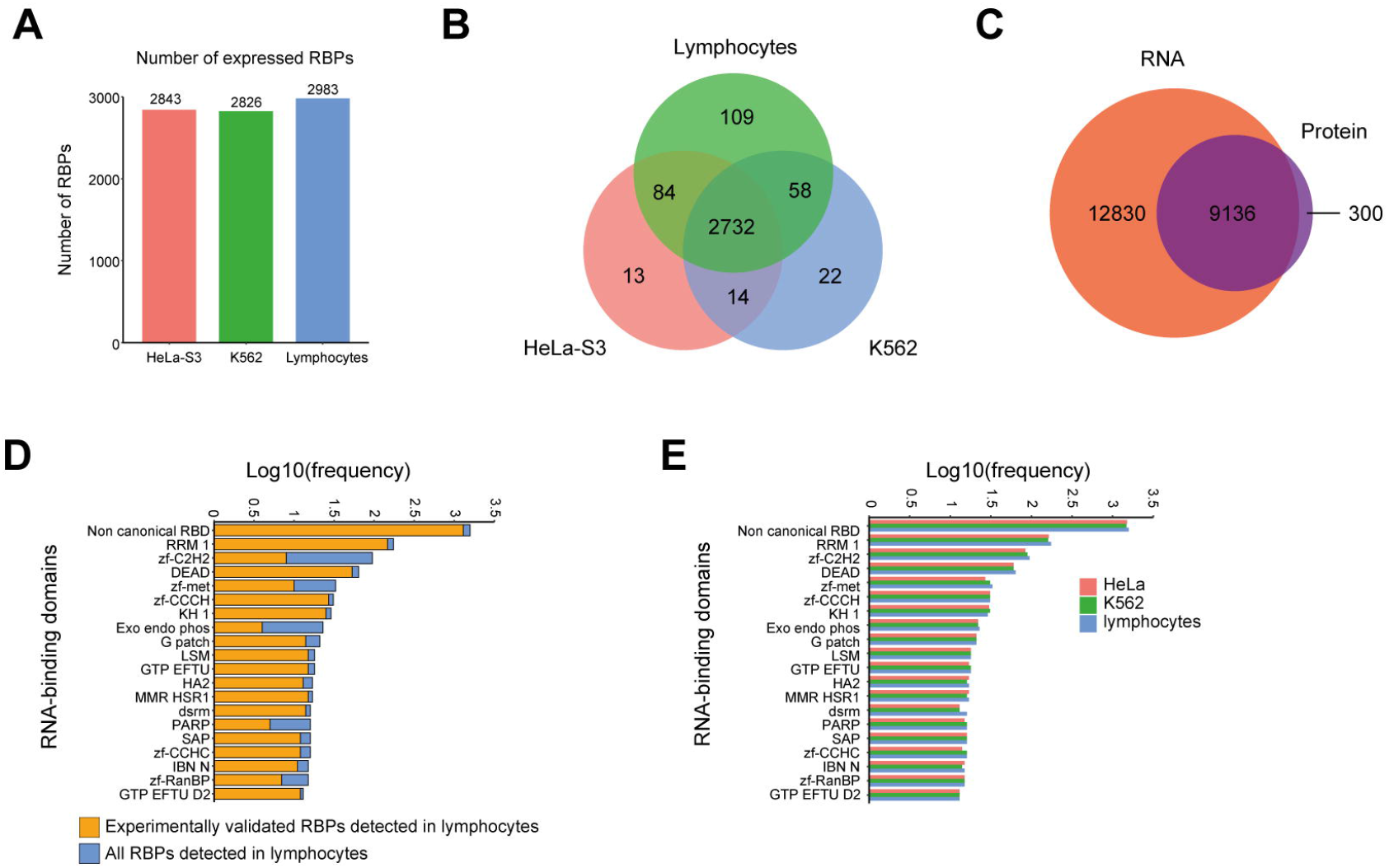

**Supplementary Figure 2.**
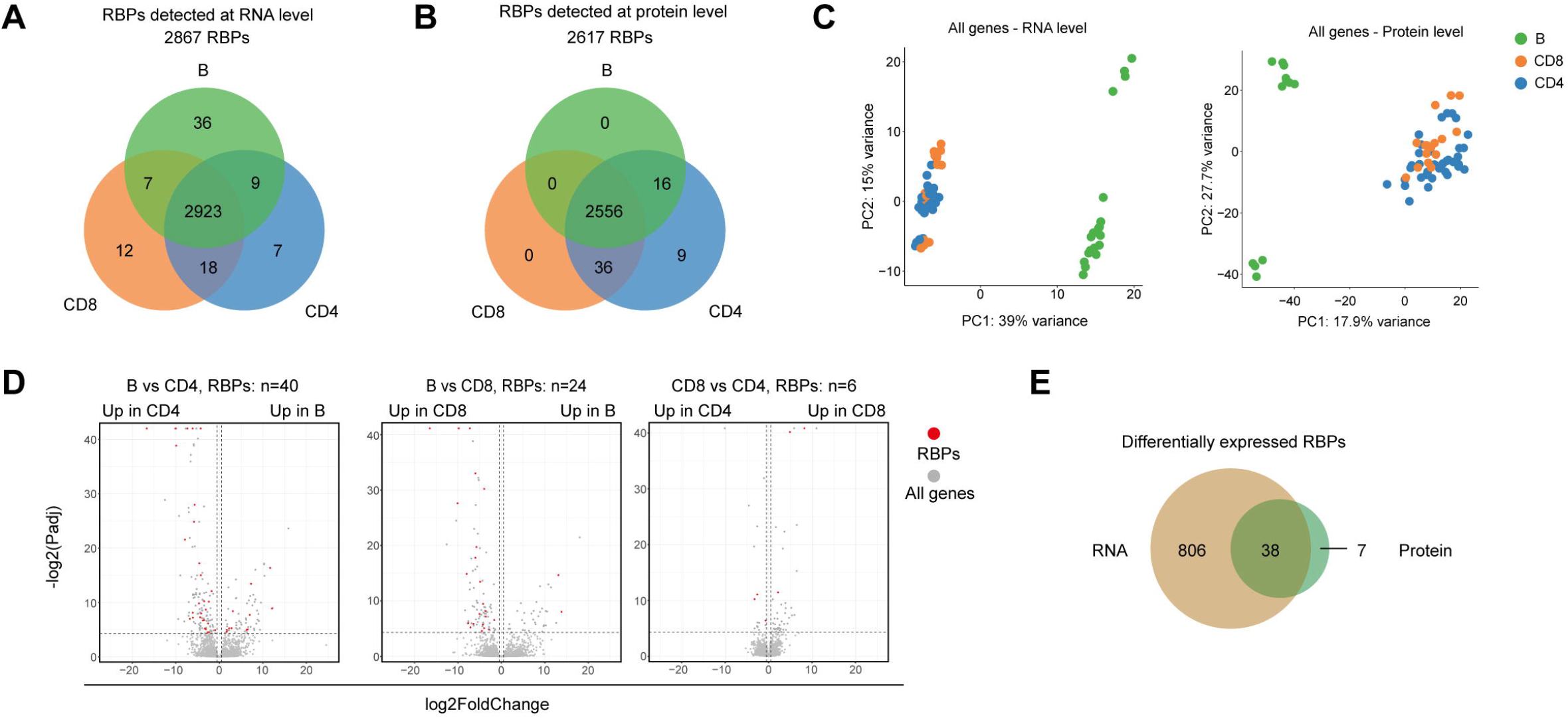

**Supplementary Figure 3.**
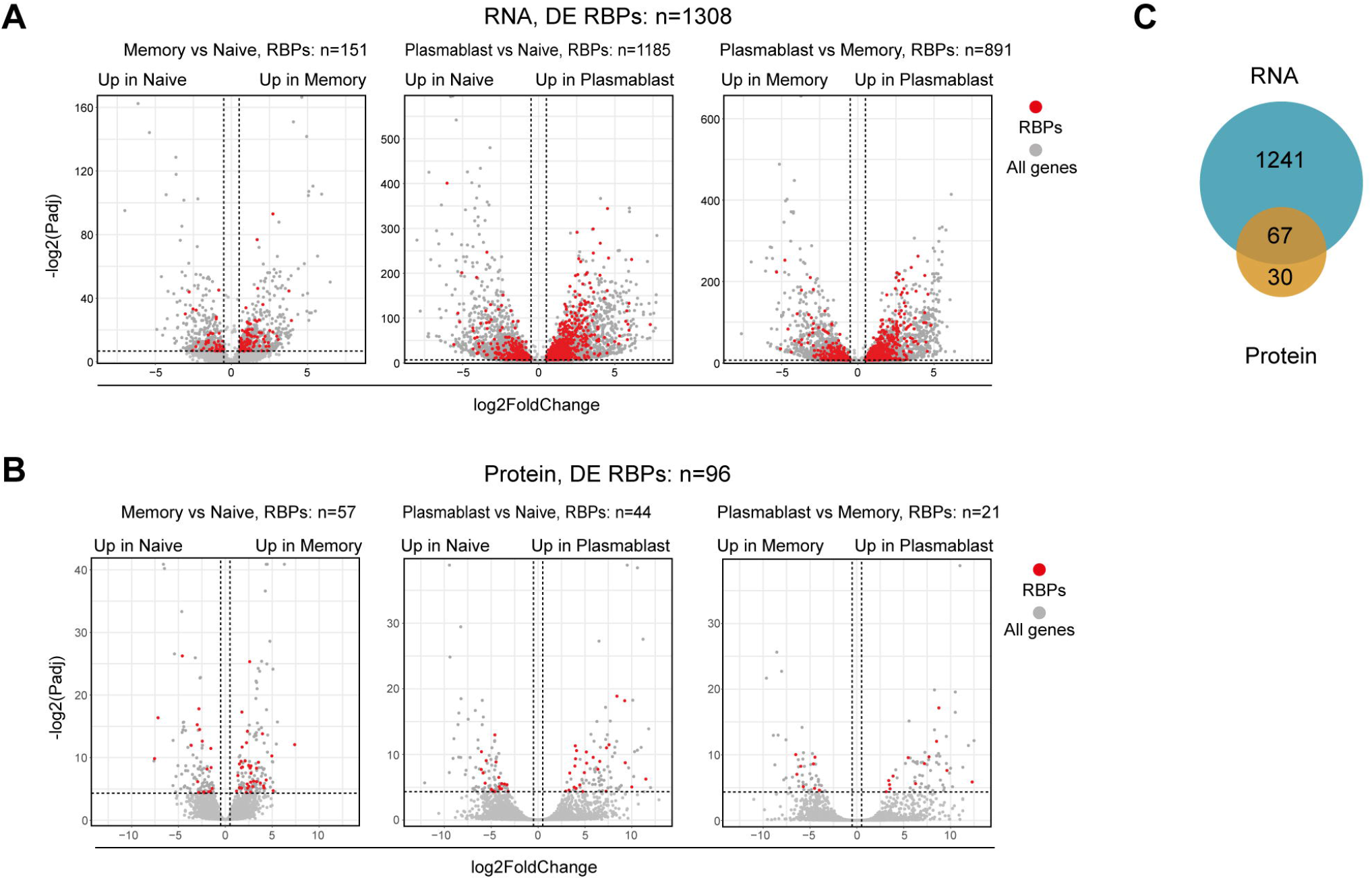

**Supplementary Figure 4.**
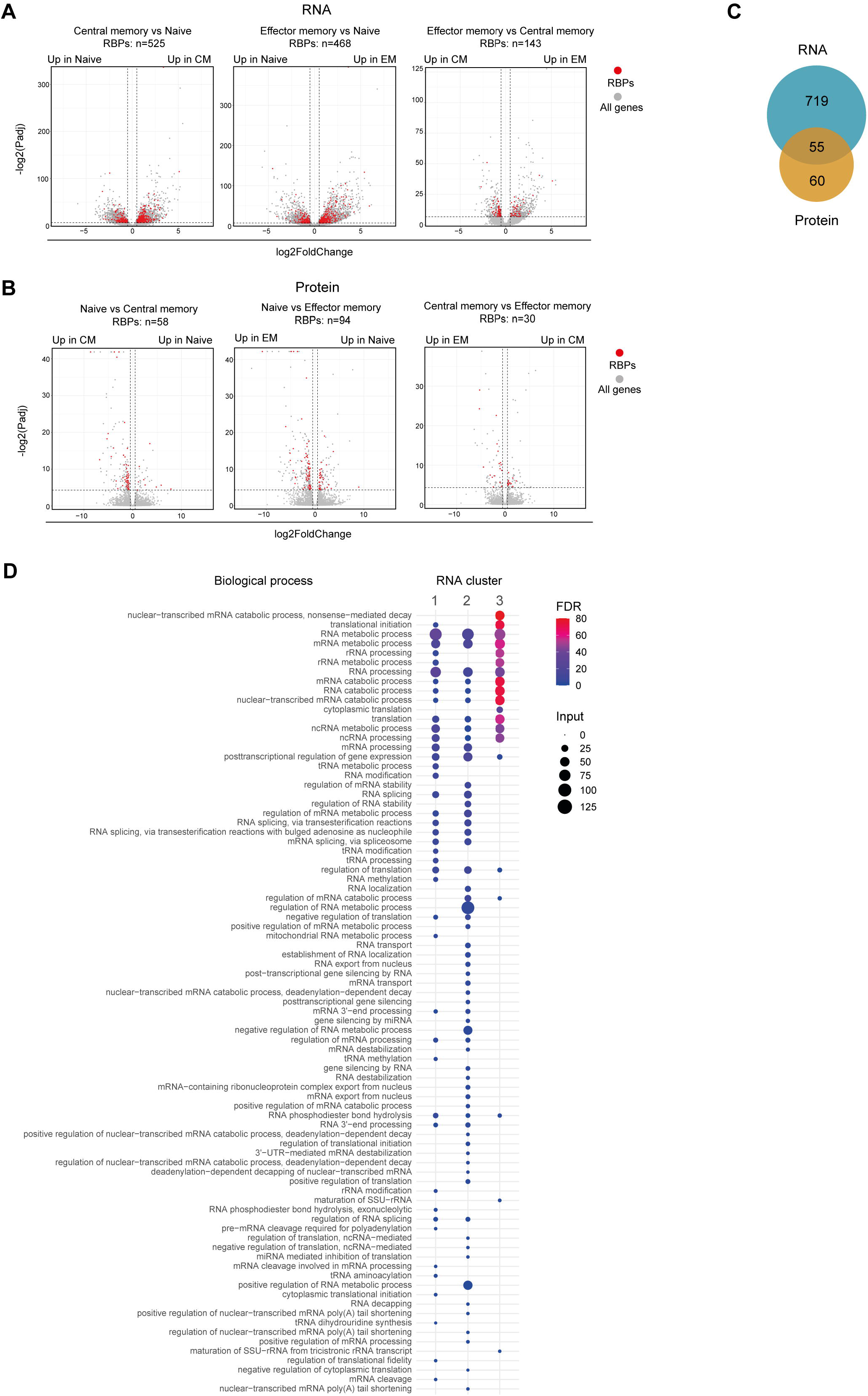

**Supplementary Figure 5.**
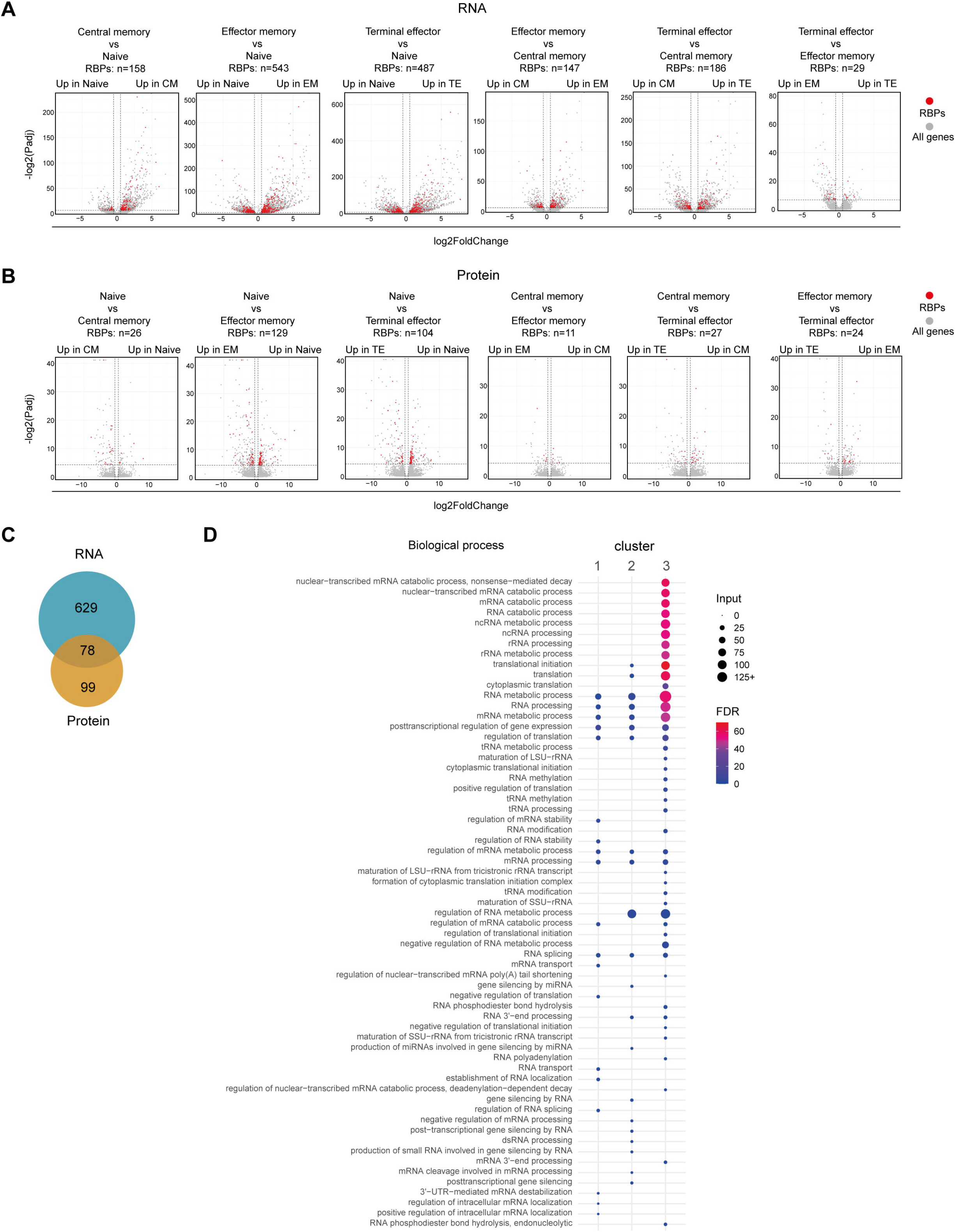

**Supp. Figure 6.**
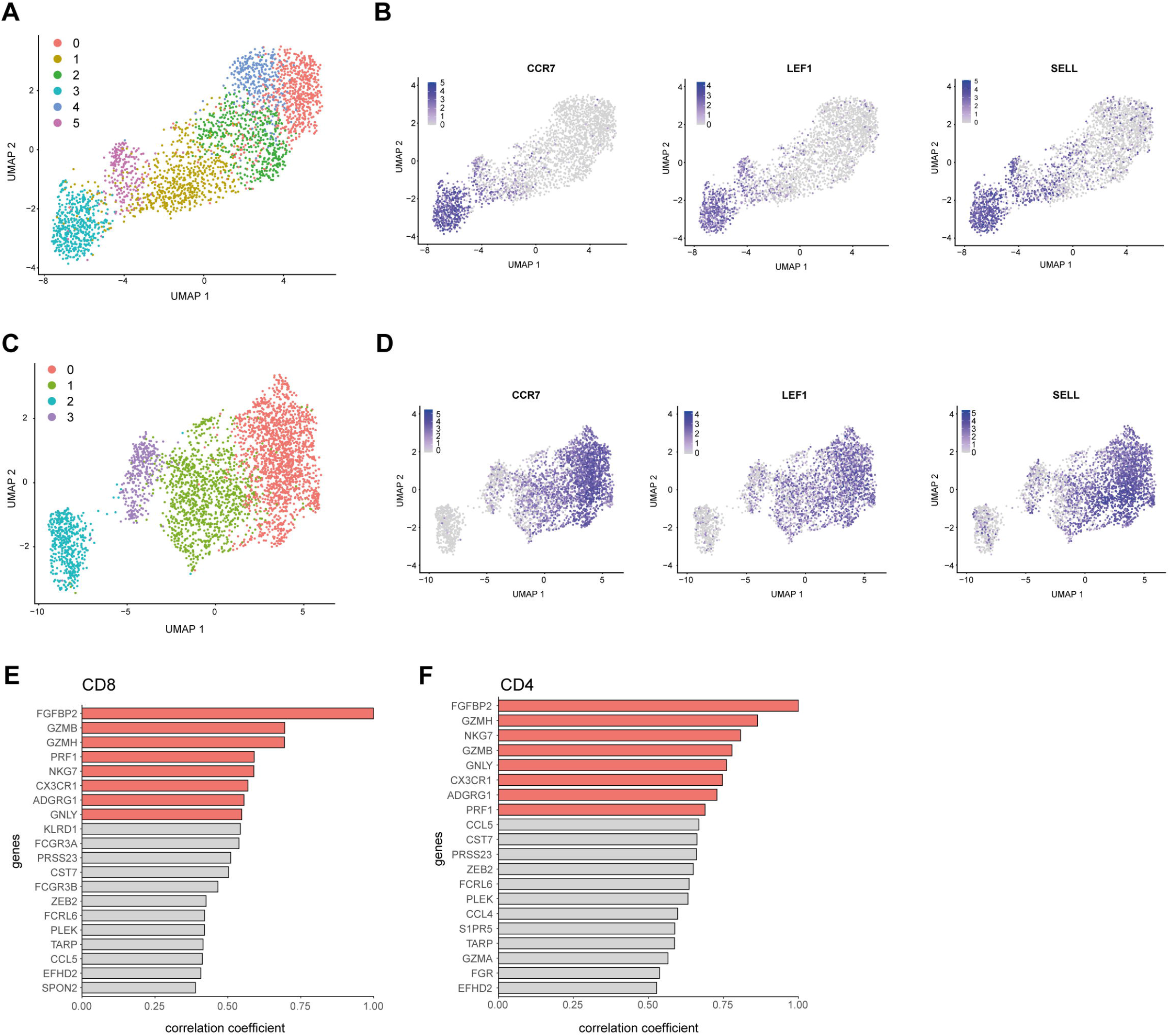

## References

1. Casanova JL, Abel L. Lethal Infectious Diseases as Inborn Errors of Immunity: Toward a Synthesis of the Germ and Genetic Theories. Annu Rev Pathol Mech Dis (2021) 16:23– 50. doi:10.1146/annurev-pathol-031920-101429

2. Dunn GP, Old LJ, Schreiber RD. The immunobiology of cancer immunosurveillance and immunoediting. Immunity (2004) 21:137–148. doi:10.1016/j.immuni.2004.07.017

3. Chemin K, Gerstner C, Malmström V. Effector functions of CD4+ T cells at the site of local autoimmune inflammation-lessons from rheumatoid arthritis. Front Immunol (2019) 10:1–15. doi:10.3389/fimmu.2019.00353

4. Das D, Akhtar S, Kurra S, Gupta S, Sharma A. Emerging role of immune cell network in autoimmune skin disorders: An update on pemphigus, vitiligo and psoriasis. Cytokine Growth Factor Rev (2019) 45:35–44. doi:10.1016/j.cytogfr.2019.01.001

5. Killestein J, Eikelenboom MJ, Izeboud T, Kalkers NF, Adèr HJ, Barkhof F, Van Lier RAW, Uitdehaag BMJ, Polman CH. Cytokine producing CD8+ T cells are correlated to MRI features of tissue destruction in MS. J Neuroimmunol (2003) 142:141–148. doi:10.1016/S0165-5728(03)00265-0

6. Arbuckle MR, McLain MT, Rubertone M V., Scofield RH, Dennis GJ, James JA. Development of auto-antibodies before the clinical onset of systemic lupus erythematosus. Rev Med Interne (2004) 25:175–176. doi:10.1016/j.revmed.2003.10.007

7. Bastard P, Rosen LB, Zhang Q, Michailidis E, Hoffmann HH, Zhang Y, Dorgham K, Philippot Q, Rosain J, Béziat V, et al. Autoantibodies against type I IFNs in patients with life-threatening COVID-19. Science (80-) (2020) 370: doi:10.1126/science.abd4585

8. Leisman DE, Ronner L, Pinotti R, Taylor MD, Sinha P, Calfee CS, Hirayama A V., Mastroiani F, Turtle CJ, Harhay MO, et al. Cytokine elevation in severe and critical COVID-19: a rapid systematic review, meta-analysis, and comparison with other inflammatory syndromes. Lancet Respir Med (2020) 8:1233–1244. doi:10.1016/S2213-2600(20)30404-5

9. Li H, van der Leun AM, Yofe I, Lubling Y, Gelbard-Solodkin D, van Akkooi ACJ, van den Braber M, Rozeman EA, Haanen JBAG, Blank CU, et al. Dysfunctional CD8 T Cells Form a Proliferative, Dynamically Regulated Compartment within Human Melanoma. Cell (2019) 176:775-789.e18. doi:10.1016/j.cell.2018.11.043

10. Fenwick C, Joo V, Jacquier P, Noto A, Banga R, Perreau M, Pantaleo G. T-cell exhaustion in HIV infection. Immunol Rev (2019) 292:149–163. doi:10.1111/imr.12823

11. Kallies A, Xin A, Belz GT, Nutt SL. Blimp-1 Transcription Factor Is Required for the Differentiation of Effector CD8+ T Cells and Memory Responses. Immunity (2009) 31:283–295. doi:10.1016/j.immuni.2009.06.021

12. Roychoudhuri R, Clever D, Li P, Wakabayashi Y, Quinn KM, Klebanoff CA, Ji Y, Sukumar M, Eil RL, Yu Z, et al. BACH2 regulates CD8 + T cell differentiation by controlling access of AP-1 factors to enhancers. Nat Immunol (2016) 17:851–860. doi:10.1038/ni.3441

13. Maciver NJ, Michalek RD, Rathmell JC. Metabolic regulation of T lymphocytes. Annu Rev Immunol (2013) 31:259–283. doi:10.1146/annurev-immunol-032712-095956

14. Boothby M, Rickert RC. Metabolic Regulation of the Immune Humoral Response. Immunity (2017) 46:743–755. doi:10.1016/j.immuni.2017.04.009

15. Rivas MA, Meydan C, Chin CR, Challman MF, Kim D, Bhinder B, Kloetgen A, Viny AD, Teater MR, McNally DR, et al. Smc3 dosage regulates B cell transit through germinal centers and restricts their malignant transformation. Nat Immunol (2021) 22:240–253. doi:10.1038/s41590-020-00827-8

16. Shin H, Blackburn SD, Intlekofer AM, Kao C, Angelosanto JM, Reiner SL, Wherry EJ. A Role for the Transcriptional Repressor Blimp-1 in CD8+ T Cell Exhaustion during Chronic Viral Infection. Immunity (2009) 31:309–320. doi:10.1016/j.immuni.2009.06.019

17. Finkin S, Hartweger H, Oliveira TY, Kara EE, Nussenzweig MC. Protein Amounts of the MYC Transcription Factor Determine Germinal Center B Cell Division Capacity. Immunity (2019) 51:324-336.e5. doi:10.1016/j.immuni.2019.06.013

18. Meffre E, Casellas R, Nussenzweig MC. Antibody regulation of B cell development. Nat Immunol (2000) 1:379–385. doi:10.1038/80816

19. Gagnon JD, Ansel KM. MicroRNA regulation of CD8+ T cell responses. Non-coding RNA Investig (2019) 3:24–24. doi:10.21037/ncri.2019.07.02

20. Salerno F, Turner M, Wolkers MC. Dynamic post-transcriptional events govern T cell homeostasis and effector function.1–31.

21. Witten JT, Ule J. Understanding splicing regulation through RNA splicing maps. Trends Genet (2011) 27:89–97. doi:10.1016/j.tig.2010.12.001

22. Müller-Mcnicoll M, Neugebauer KM. How cells get the message: Dynamic assembly and function of mRNA-protein complexes. Nat Rev Genet (2013) 14:275–287. doi:10.1038/nrg3434

23. Song J, Yi C. Chemical Modifications to RNA: A New Layer of Gene Expression Regulation. ACS Chem Biol (2017) 12:316–325. doi:10.1021/acschembio.6b00960

24. Alison Galloway, Alexander Saveliev, Sebastian Łukasiak, Daniel J. Hodson, Daniel Bolland, Kathryn Balmanno, Helena Ahlfors, 1 Elisa Monzón-Casanova, Sara Ciullini Mannurita, Lewis S. Bell, Simon Andrews, Manuel D. Díaz-Muñoz, Simon J. Cook, Anne Corcoran MT. RNA-binding proteins ZFP36L1 and ZFP36L2 promote cell quiescence. Science (80-) (2016) 352:453–459. doi:10.1126/science.aad5978

25. Newman R, Ahlfors H, Saveliev A, Galloway A, Hodson DJ, Williams R, Besra GS, Cook CN, Cunningham AF, Bell SE, et al. Maintenance of the marginal-zone B cell compartment specifically requires the RNA-binding protein ZFP36L1. Nat Immunol (2017) 18:683–693. doi:10.1038/ni.3724

26. Monzón-Casanova E, Screen M, Díaz-Muñoz MD, Coulson RMR, Bell SE, Lamers G, Solimena M, Smith CWJ, Turner M. The RNA-binding protein PTBP1 is necessary for B cell selection in germinal centers article. Nat Immunol (2018) 19:267–278. doi:10.1038/s41590-017-0035-5

27. Galloway A, Ahlfors H, Turner M, Bell LS, Vogel KU. The RNA-Binding Proteins Zfp36l1 and Zfp36l2 Enforce the Thymic β-Selection Checkpoint by Limiting DNA Damage Response Signaling and Cell Cycle Progression. J Immunol (2016) 197:2673– 2685. doi:10.4049/jimmunol.1600854

28. Essig K, Hu D, Guimaraes JC, Alterauge D, Edelmann S, Raj T, Kranich J, Behrens G, Heiseke A, Floess S, et al. Roquin Suppresses the PI3K-mTOR Signaling Pathway to Inhibit T Helper Cell Differentiation and Conversion of Treg to Tfr Cells. Immunity (2017) 47:1067-1082.e12. doi:10.1016/j.immuni.2017.11.008

29. Cui X, Mino T, Yoshinaga M, Nakatsuka Y, Hia F, Yamasoba D, Tsujimura T, Tomonaga K, Suzuki Y, Uehata T, et al. Regnase-1 and Roquin Nonredundantly Regulate Th1 Differentiation Causing Cardiac Inflammation and Fibrosis. J Immunol (2017) 199:4066–4077. doi:10.4049/jimmunol.1701211

30. Li HB, Tong J, Zhu S, Batista PJ, Duffy EE, Zhao J, Bailis W, Cao G, Kroehling L, Chen Y, et al. M 6 A mRNA methylation controls T cell homeostasis by targeting the IL-7/STAT5/SOCS pathways. Nature (2017) 548:338–342. doi:10.1038/nature23450

31. Wei J, Long L, Zheng W, Dhungana Y, Lim SA, Guy C, Wang Y, Wang YD, Qian C, Xu B, et al. Targeting REGNASE-1 programs long-lived effector T cells for cancer therapy. Nature (2019) 576:471–476. doi:10.1038/s41586-019-1821-z

32. Tavernier SJ, Athanasopoulos V, Verloo P, Behrens G, Staal J, Bogaert DJ, Naesens L, De Bruyne M, Van Gassen S, Parthoens E, et al. A human immune dysregulation syndrome characterized by severe hyperinflammation with a homozygous nonsense Roquin-1 mutation. Nat Commun (2019) 10: doi:10.1038/s41467-019-12704-6

33. Salerno F, Engels S, van den Biggelaar M, van Alphen FPJ, Guislain A, Zhao W, Hodge DL, Bell SE, Medema JP, von Lindern M, et al. Translational repression of pre-formed cytokine-encoding mRNA prevents chronic activation of memory T cells. Nat Immunol (2018) 19:828–837. doi:10.1038/s41590-018-0155-6

34. Castello A, Fischer B, Frese CK, Horos R, Alleaume AM, Foehr S, Curk T, Krijgsveld J, Hentze MW. Comprehensive Identification of RNA-Binding Domains in Human Cells. Mol Cell (2016) 63:696–710. doi:10.1016/j.molcel.2016.06.029

35. Perez-Perri JI, Rogell B, Schwarzl T, Stein F, Zhou Y, Rettel M, Brosig A, Hentze MW. Discovery of RNA-binding proteins and characterization of their dynamic responses by enhanced RNA interactome capture. Nat Commun (2018) 9: doi:10.1038/s41467-018-06557-8

36. Liao JY, Yang B, Zhang YC, Wang XJ, Ye Y, Peng JW, Yang ZZ, He JH, Zhang Y, Hu KS, et al. EuRBPDB: a comprehensive resource for annotation, functional and oncological investigation of eukaryotic RNA binding proteins (RBPs). Nucleic Acids Res (2020) 48:D307–D313. doi:10.1093/nar/gkz823

37. Gerstberger S, Hafner M, Tuschl T. A census of human RNA-binding proteins. Nat Rev Genet (2014) 15:829–845. doi:10.1038/nrg3813

38. Monaco G, Lee B, Xu W, Mustafah S, Hwang YY, Carré C, Burdin N, Visan L, Ceccarelli M, Poidinger M, et al. RNA-Seq Signatures Normalized by mRNA Abundance Allow Absolute Deconvolution of Human Immune Cell Types. Cell Rep (2019) 26:1627-1640.e7. doi:10.1016/j.celrep.2019.01.041

39. Martinez TF, Chu Q, Donaldson C, Tan D, Shokhirev MN, Saghatelian A. Accurate annotation of human protein-coding small open reading frames. Nat Chem Biol (2020) 16:458–468. doi:10.1038/s41589-019-0425-0

40. Rieckmann JC, Geiger R, Hornburg D, Wolf T, Kveler K, Jarrossay D, Sallusto F, Shen-Orr SS, Lanzavecchia A, Mann M, et al. Social network architecture of human immune cells unveiled by quantitative proteomics. Nat Immunol (2017) 18:583–593. doi:10.1038/ni.3693

41. Legrand JMD, Chan AL, La HM, Rossello FJ, Änkö ML, Fuller-Pace F V., Hobbs RM. DDX5 plays essential transcriptional and post-transcriptional roles in the maintenance and function of spermatogonia. Nat Commun (2019) 10: doi:10.1038/s41467-019-09972-7

42. Diamond RH, D. K, Lee VM, Mohn KL, Haber BA, Tewari DS, Taub R. Novel delayed-early and highly insulin-induced growth response genes. Identification of HRS, a potential regulator of alternative pre-mRNA splicing. J Biol Chem (1993) 268:15185– 15192. doi:10.1016/s0021-9258(18)82454-1

43. Mayeda A, Krainer AR. Regulation of alternative pre-mRNA splicing by hnRNP A1 and splicing factor SF2. Cell (1992) 68:365–375. doi:10.1016/0092-8674(92)90477-T

44. Challa AA, Stefanovic B. A Novel Role of Vimentin Filaments: Binding and Stabilization of Collagen mRNAs. Mol Cell Biol (2011) 31:3773–3789. doi:10.1128/mcb.05263-11

45. Alarcón CR, Goodarzi H, Lee H, Liu X, Tavazoie S, Tavazoie SF. HNRNPA2B1 Is a Mediator of m6A-Dependent Nuclear RNA Processing Events. Cell (2015) 162:1299– 1308. doi:10.1016/j.cell.2015.08.011

46. Castello A, Hentze MW, Preiss T. Metabolic Enzymes Enjoying New Partnerships as RNA-Binding Proteins. Trends Endocrinol Metab (2015) 26:746–757. doi:10.1016/j.tem.2015.09.012

47. Chang CH, Curtis JD, Maggi LB, Faubert B, Villarino A V., O’Sullivan D, Huang SCC, Van Der Windt GJW, Blagih J, Qiu J, et al. Posttranscriptional control of T cell effector function by aerobic glycolysis. Cell (2013) 153:1239. doi:10.1016/j.cell.2013.05.016

48. Finn RD, Bateman A, Clements J, Coggill P, Eberhardt RY, Eddy SR, Heger A, Hetherington K, Holm L, Mistry J, et al. Pfam: The protein families database. Nucleic Acids Res (2014) 42:222–230. doi:10.1093/nar/gkt1223

49. Díaz-Muñoz MD, Turner M. Uncovering the role of RNA-binding proteins in gene expression in the immune system. Front Immunol (2018) 9: doi:10.3389/fimmu.2018.01094

50. Uhlén M, Fagerberg L, Hallström BM, Lindskog C, Oksvold P, Mardinoglu A, Sivertsson Å, Kampf C, Sjöstedt E, Asplund A, et al. Tissue-based map of the human proteome. Science (80-) (2015) 347: doi:10.1126/science.1260419

51. Nutt SL, Hodgkin PD, Tarlinton DM, Corcoran LM. The generation of antibody-secreting plasma cells. Nat Rev Immunol (2015) 15:160–171. doi:10.1038/nri3795

52. Battle DJ, Doudna JA. The stem-loop binding protein forms a highly stable and specific complex with the 3′ stem-loop of histone mRNAs. Rna (2001) 7:123–132. doi:10.1017/S1355838201001820

53. Smirnov A, Comte C, Mager-Heckel AM, Addis V, Krasheninnikov IA, Martin RP, Entelis N, Tarassov I. Mitochondrial enzyme rhodanese is essential for 5 S ribosomal RNA import into human mitochondria. J Biol Chem (2010) 285:30792–30803. doi:10.1074/jbc.M110.151183

54. Ding Y, Zhou L, Xia Y, Wang W, Wang Y, Li L, Qi Z, Zhong L, Sun J, Tang W, et al. Reference values for peripheral blood lymphocyte subsets of healthy children in China. J Allergy Clin Immunol (2018) 142:970-973.e8. doi:10.1016/j.jaci.2018.04.022

55. Sallusto F, Lenig D, Förster R, Lipp M, Lanzavecchia A. Two subsets of memory T lymphocytes with distinct homing potentials and effector functions. Nature (1999) 401:708–712. doi:10.1038/44385

56. Tran DDH, Koch A, Tamura T. THOC5, a member of the mRNA export complex: A novel link between mRNA export machinery and signal transduction pathways in cell proliferation and differentiation. Cell Commun Signal (2014) 12:1–9. doi:10.1186/1478-811X-12-3

57. Chuang TW, Peng PJ, Tarn WY. The exon junction complex component Y14 modulates the activity of the methylosome in biogenesis of spliceosomal small nuclear ribonucleoproteins. J Biol Chem (2011) 286:8722–8728. doi:10.1074/jbc.M110.190587

58. Dufu K, Livingstone MJ, Seebacher J, Gygi SP, Wilson SA, Reed R. ATP is required for interactions between UAP56 and two conserved mRNA export proteins, Aly and CIP29, to assemble the TREX complex. Genes Dev (2010) 24:2043–2053. doi:10.1101/gad.1898610

59. Nicolet BP, Guislain A, van Alphen FPJ, Gomez-Eerland R, Schumacher TNM, van den Biggelaar M, Wolkers MC. CD29 identifies IFN-γ-producing human CD8+ T cells with an increased cytotoxic potential. Proc Natl Acad Sci U S A (2020) 117:6686–6696. doi:10.1073/pnas.1913940117

60. St. Paul M, Ohashi PS. The Roles of CD8+ T Cell Subsets in Antitumor Immunity. Trends Cell Biol (2020) 30:695–704. doi:10.1016/j.tcb.2020.06.003

61. Loyal L, Warth S, Jürchott K, Mölder F, Nikolaou C, Babel N, Nienen M, Durlanik S, Stark R, Kruse B, et al. SLAMF7 and IL-6R define distinct cytotoxic versus helper memory CD8+ T cells. Nat Commun (2020) 11: doi:10.1038/s41467-020-19002-6

62. Nicolet BP, Guislain A, Wolkers MC. CD29 enriches for cytotoxic human CD4 T cells. (2021) doi:https://doi.org/10.1101/2021.02.10.430576

63. Oh DY, Kwek SS, Raju SS, Li T, McCarthy E, Chow E, Aran D, Ilano A, Pai CCS, Rancan C, et al. Intratumoral CD4+ T Cells Mediate Anti-tumor Cytotoxicity in Human Bladder Cancer. Cell (2020) 181:1612-1625.e13. doi:10.1016/j.cell.2020.05.017

64. Cachot A, Bilous M, Liu Y-C, Li X, Rockinger A, Saillard M, Wyss T, Guillaume P, Schmidt J, Genolet R, et al. Tumor-specific cytolytic CD4 T cells mediate protective immunity against human cancer. J Immunother Cancer (2020) 8:A581–A581. doi:10.1136/jitc-2020-sitc2020.0545

65. Guo X, Zhang Y, Zheng L, Zheng C, Song J, Zhang Q, Kang B, Liu Z, Jin L, Xing R, et al. Global characterization of T cells in non-small-cell lung cancer by single-cell sequencing. Nat Med (2018) 24:978–985. doi:10.1038/s41591-018-0045-3

66. Zhang Y, Zheng L, Zhang L, Hu X, Ren X, Zhang Z. Deep single-cell RNA sequencing data of individual T cells from treatment-naïve colorectal cancer patients. Sci data (2019) 6:131. doi:10.1038/s41597-019-0131-5

67. Zheng C, Zheng L, Yoo JK, Guo H, Zhang Y, Guo X, Kang B, Hu R, Huang JY, Zhang Q, et al. Landscape of Infiltrating T Cells in Liver Cancer Revealed by Single-Cell Sequencing. Cell (2017) 169:1342-1356.e16. doi:10.1016/j.cell.2017.05.035

68. Shi Z, Fujii K, Kovary KM, Genuth NR, Röst HL, Teruel MN, Barna M. Heterogeneous Ribosomes Preferentially Translate Distinct Subpools of mRNAs Genome-wide. Mol Cell (2017) 67:71-83.e7. doi:10.1016/j.molcel.2017.05.021

69. Filipenko NR, MacLeod TJ, Yoon CS, Waisman DM. Annexin A2 Is a Novel RNA-binding Protein. J Biol Chem (2004) 279:8723–8731. doi:10.1074/jbc.M311951200

70. Ishibashi M, Wakita T, Esumi M. 2′,5′-Oligoadenylate synthetase-like gene highly induced by hepatitis C virus infection in human liver is inhibitory to viral replication in vitro. Biochem Biophys Res Commun (2010) 392:397–402. doi:10.1016/j.bbrc.2010.01.034

71. Marnef A, Maldonado M, Bugaut A, Balasubramanian S, Kress M, Weil D, Standart N. Distinct functions of maternal and somatic Pat1 protein paralogs. Rna (2010) 16:2094– 2107. doi:10.1261/rna.2295410

72. Wolf T, Jin W, Zoppi G, Vogel IA, Akhmedov M, Bleck CKE, Beltraminelli T, Rieckmann JC, Ramirez NJ, Benevento M, et al. Dynamics in protein translation sustaining T cell preparedness. Nat Immunol (2020) 21:927–937. doi:10.1038/s41590-020-0714-5

73. Howden AJM, Hukelmann JL, Brenes A, Spinelli L, Sinclair L V., Lamond AI, Cantrell DA. Quantitative analysis of T cell proteomes and environmental sensors during T cell differentiation. Nat Immunol (2019) 20:1542–1554. doi:10.1038/s41590-019-0495-x

74. Myers DR, Norlin E, Vercoulen Y, Roose JP. Active Tonic mTORC1 Signals Shape Baseline Translation in Naive T Cells. Cell Rep (2019) 27:1858-1874.e6. doi:10.1016/j.celrep.2019.04.037

75. Araki K, Morita M, Bederman AG, Konieczny BT, Kissick HT, Sonenberg N, Ahmed R. Translation is actively regulated during the differentiation of CD8 + effector T cells. Nat Immunol (2017) 18:1046–1057. doi:10.1038/ni.3795

76. Refsland EW, Stenglein MD, Shindo K, Albin JS, Brown WL, Harris RS. Quantitative profiling of the full APOBEC3 mRNA repertoire in lymphocytes and tissues: Implications for HIV-1 restriction. Nucleic Acids Res (2010) 38:4274–4284. doi:10.1093/nar/gkq174

77. Gillick K, Pollpeter D, Phalora P, Kim E-Y, Wolinsky SM, Malim MH. Suppression of HIV-1 Infection by APOBEC3 Proteins in Primary Human CD4 + T Cells Is Associated with Inhibition of Processive Reverse Transcription as Well as Excessive Cytidine Deamination . J Virol (2013) 87:1508–1517. doi:10.1128/jvi.02587-12

78. Beckmann BM, Horos R, Fischer B, Castello A, Eichelbaum K, Alleaume AM, Schwarzl T, Curk T, Foehr S, Huber W, et al. The RNA-binding proteomes from yeast to man harbour conserved enigmRBPs. Nat Commun (2015) 6: doi:10.1038/ncomms10127

79. Huppertz I, Perez-Perri JI, Mantas P, Sekaran T, Schwarzl T, Dimitrova-Paternoga L, Hennig J, Neveu PA, Hentze MW. RNA regulates Glycolysis and Embryonic Stem Cell Differentiation via Enolase 1. bioRxiv (2020)2020.10.14.337444. Available at: https://doi.org/10.1101/2020.10.14.337444

80. Gebauer F, Schwarzl T, Valcárcel J, Hentze MW. RNA-binding proteins in human genetic disease. Nat Rev Genet (2021) 22:185–198. doi:10.1038/s41576-020-00302-y

81. Ranzani V, Rossetti G, Panzeri I, Arrigoni A, Bonnal RJP, Curti S, Gruarin P, Provasi E, Sugliano E, Marconi M, et al. The long intergenic noncoding RNA landscape of human lymphocytes highlights the regulation of T cell differentiation by linc-MAF-4. Nat Immunol (2015) 16:318–325. doi:10.1038/ni.3093

82. Zhang L, Yu X, Zheng L, Zhang Y, Li Y, Fang Q, Gao R, Kang B, Zhang Q, Huang JY, et al. Lineage tracking reveals dynamic relationships of T cells in colorectal cancer. Nature (2018) 564:268–272. doi:10.1038/s41586-018-0694-x

83. Patro R, Duggal G, Love MI, Irizarry RA, Kingsford C. Salmon provides fast and bias-aware quantification of transcript expression. Nat Methods (2017) 14:417–419. doi:10.1038/nmeth.4197

84. Soneson C, Love MI, Robinson MD. Differential analyses for RNA-seq: Transcript-level estimates improve gene-level inferences [version 2; referees: 2 approved]. F1000Research (2016) 4:1–23. doi:10.12688/F1000RESEARCH.7563.2

85. Love MI, Huber W, Anders S. Moderated estimation of fold change and dispersion for RNA-seq data with DESeq2. Genome Biol (2014) 15:1–21. doi:10.1186/s13059-014-0550-8

86. Satija R, Farrell JA, Gennert D, Schier AF, Regev A. Spatial reconstruction of single-cell gene expression data. Nat Biotechnol (2015) 33:495–502. doi:10.1038/nbt.3192

87. Stuart T, Butler A, Hoffman P, Hafemeister C, Papalexi E, Mauck WM, Hao Y, Stoeckius M, Smibert P, Satija R. Comprehensive Integration of Single-Cell Data. Cell (2019) 177:1888-1902.e21. doi:10.1016/j.cell.2019.05.031

88. Willinger T, Freeman T, Herbert M, Hasegawa H, McMichael AJ, Callan MFC. Human Naive CD8 T Cells Down-Regulate Expression of the WNT Pathway Transcription Factors Lymphoid Enhancer Binding Factor 1 and Transcription Factor 7 (T Cell Factor-1) following Antigen Encounter In Vitro and In Vivo. J Immunol (2006) 176:1439–1446. doi:10.4049/jimmunol.176.3.1439

89. Yang C, Khanniche A, Dispirito JR, Ji P, Wang S, Wang Y, Shen H. Transcriptome Signatures Reveal Rapid Induction of Immune-Responsive Genes in Human Memory CD8+ T Cells. Sci Rep (2016) 6:1–8. doi:10.1038/srep27005

90. Finak G, McDavid A, Yajima M, Deng J, Gersuk V, Shalek AK, Slichter CK, Miller HW, McElrath MJ, Prlic M, et al. MAST: A flexible statistical framework for assessing transcriptional changes and characterizing heterogeneity in single-cell RNA sequencing data. Genome Biol (2015) 16:1–13. doi:10.1186/s13059-015-0844-5

91. Zhang X, Smits AH, Van Tilburg GBA, Ovaa H, Huber W, Vermeulen M. Proteome-wide identification of ubiquitin interactions using UbIA-MS. Nat Protoc (2018) 13:530– 550. doi:10.1038/nprot.2017.147

92. Szklarczyk D, Gable AL, Lyon D, Junge A, Wyder S, Huerta-Cepas J, Simonovic M, Doncheva NT, Morris JH, Bork P, et al. STRING v11: Protein-protein association networks with increased coverage, supporting functional discovery in genome-wide experimental datasets. Nucleic Acids Res (2019) 47:D607–D613. doi:10.1093/nar/gky1131

93. Mi H, Muruganujan A, Ebert D, Huang X, Thomas PD. PANTHER version 14: More genomes, a new PANTHER GO-slim and improvements in enrichment analysis tools. Nucleic Acids Res (2019) 47:D419–D426. doi:10.1093/nar/gky1038

94. Wickham H. ggplot2: Elegant Graphics for Data Analysis. Springer-Verlag (2016) ISBN 978-3:

95. Kolde R. pheatmap: Pretty heatmaps. URL https://CRANR-projectorg/package=pheatmap (2015)

