## Supplementary Figures for "Mapping RNA-binding proteins in human B cells and T cells upon differentiation"

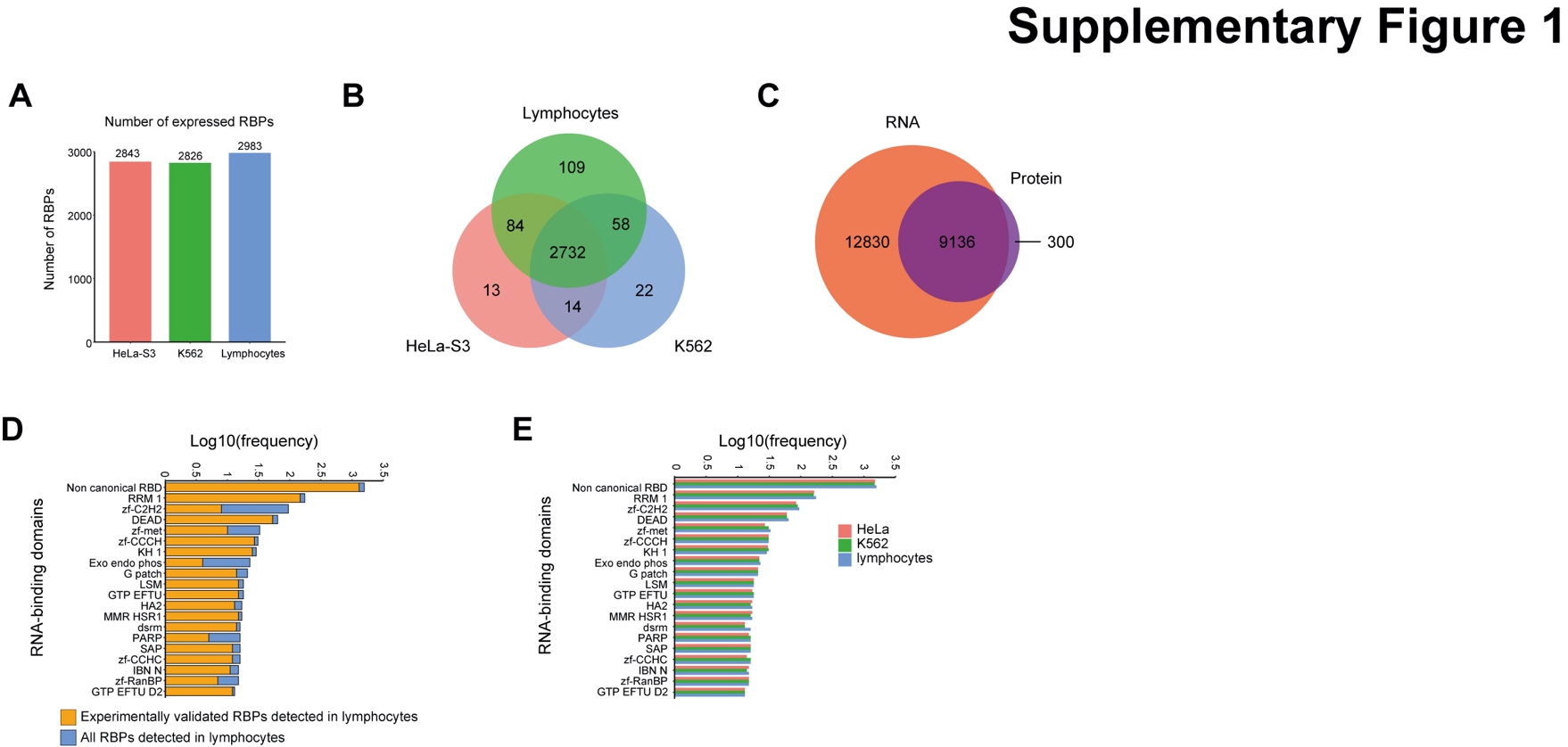

**Supplementary Figure 1:** **Characterization of RBP expression in human B and T lymphocytes.**

(**A**) Number of RBPs detected at RNA level in HeLa-S3 and K562 cells compared to human B/T lymphocytes. (**B**) Venn diagram depicting the number of RBP genes identified in indicated cell type. (**C**) Venn diagram depicting the number of gene products detected at RNA (orange) and protein level (purple) in human B/T lymphocytes. (**D**) Bar chart depicting the frequency of RNA-binding domains (RBDs) in B/T lymphocytes among all identified RBPs (blue) and among experimentally validated RBPs (orange). (**E**) Bar chart depicting the frequency of RBDs in RBPs detected in indicated cell types.

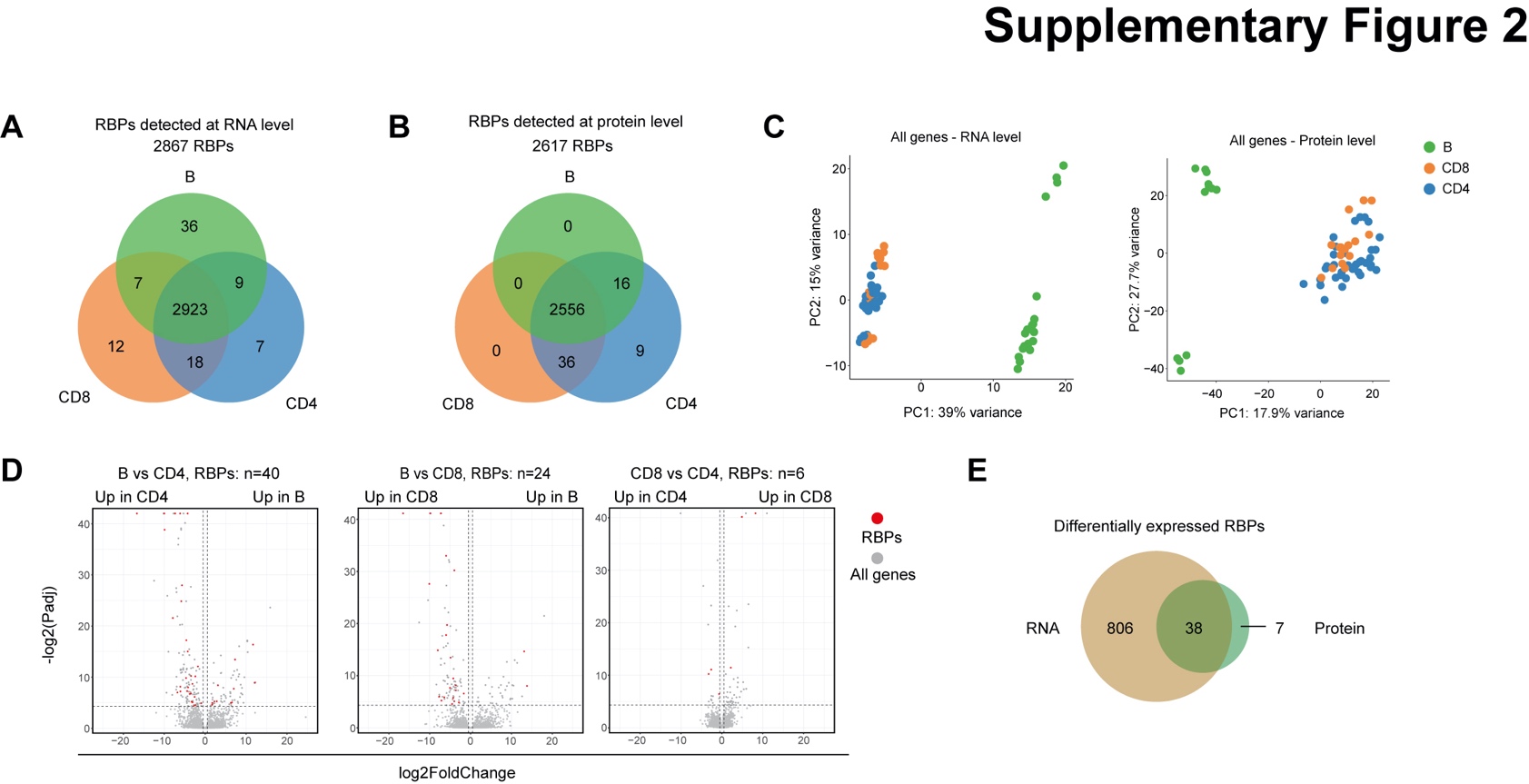

**Supplementary Figure 2:** **Differential RBP expression in human B cells and T cells.**

(**A**-**B**) RBPs detected at RNA (**A**) and protein level (**B**) in human CD19^+^ B cells, CD4^+^ T cells and CD8^+^ T cells. (**C**) Left panel: Principal component analysis (PCA) plot of RNA (left panel) and protein expression (right panel) of all gene products in CD19^+^ B cells. Each dot depicts one subset from 3-4 donors. (**D**) Volcano plots depicting differential expression of all proteins (gray), and of RBPs (red) between lymphocyte subsets (LFC>0.5, P-adjusted<0.05). (**E**) Venn diagram showing the numbers and overlay between DE RBPs at RNA and protein level.

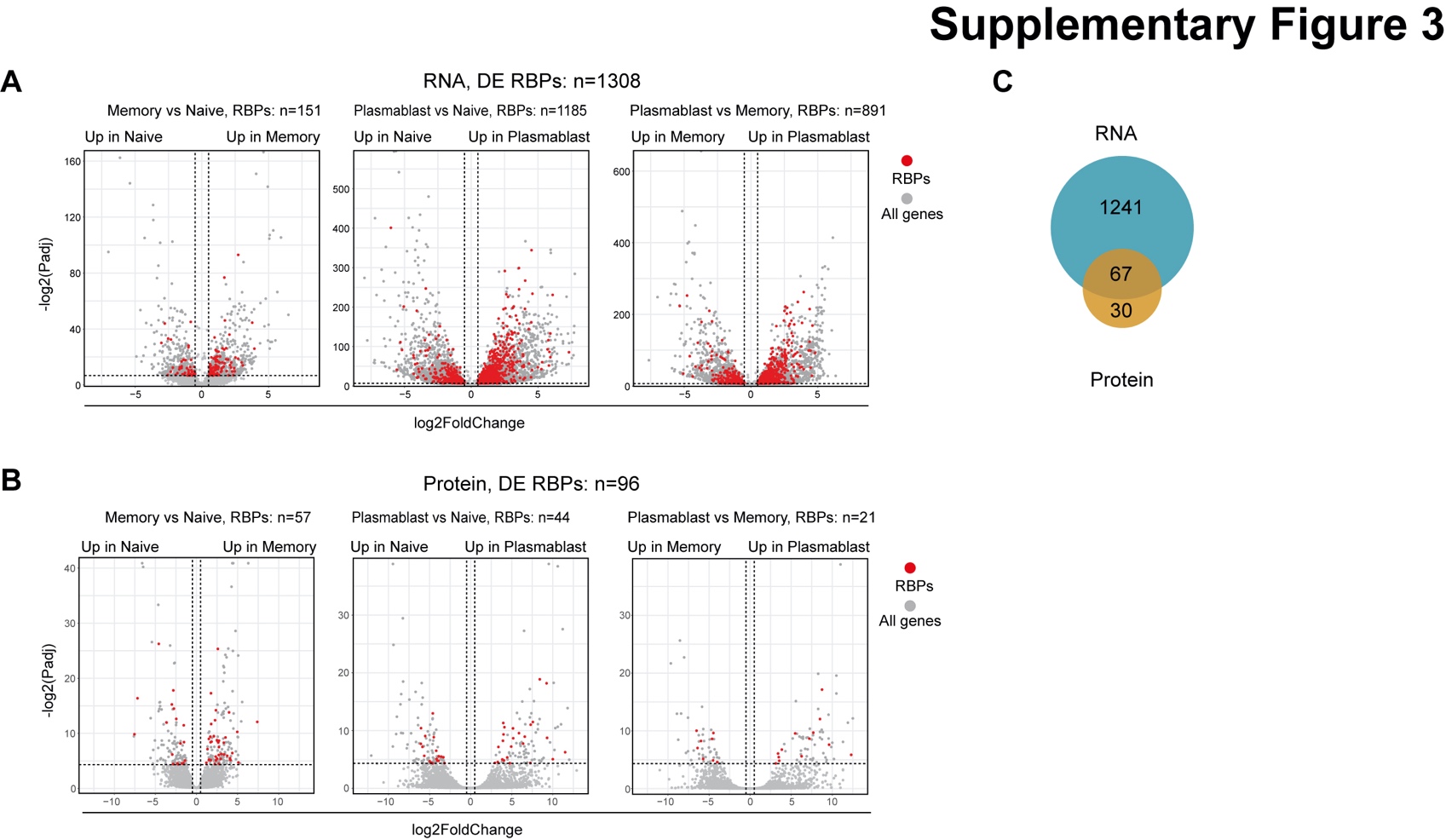

**Supplementary Figure 3:** **Differential RBP expression in B cell subsets.**

(**A-B**) Volcano plots depicting differential RNA (LFC>0.5, P-adjusted<0.01) **(A**) and protein expression (LFC>0.5, P-adjusted<0.05 **(B)** of all genes (gray) and of RBPs (red) between human CD19^+^ B cell subsets. (**C**) Venn diagram showing the overlay between RBP RNA and protein expression in CD19^+^ B cells.

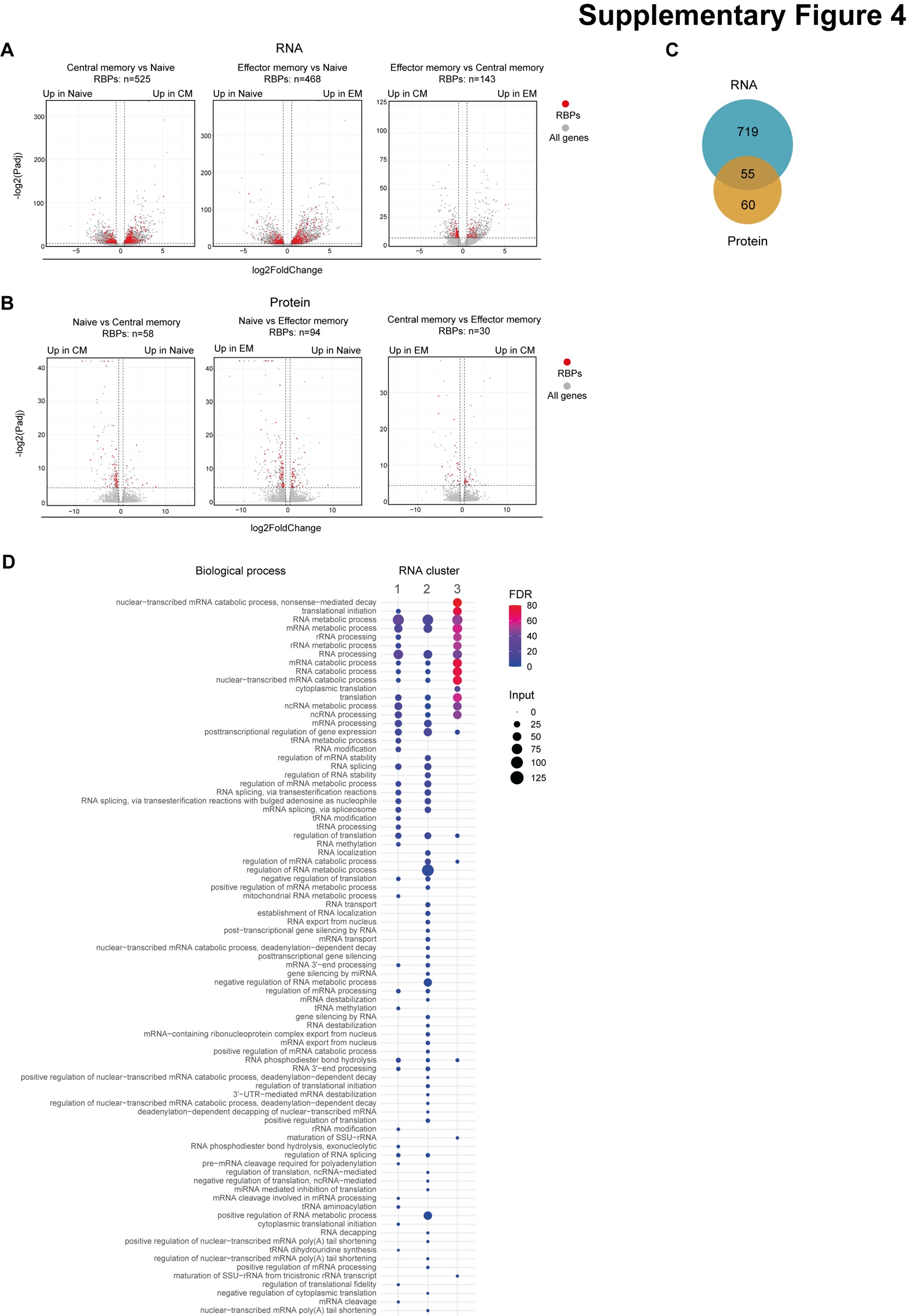

**Supplementary Figure 4:** **Differential RBP expression and GO-term analysis of RBPs in human CD4^+^**

**T cells.**

(**A-B**) Volcano plots depicting differential RNA **(A**) and protein expression **(B)** of all genes (gray) and of RBPs (red) between human CD4^+^ T cell subsets (LFC>0.5, P-adjusted<0.01). (**C**) Venn diagram showing the overlay between RBP RNA and protein expression. (**D**) RNA-related biological processes enriched in clusters identified in Figure 4B (FDR<0.05).

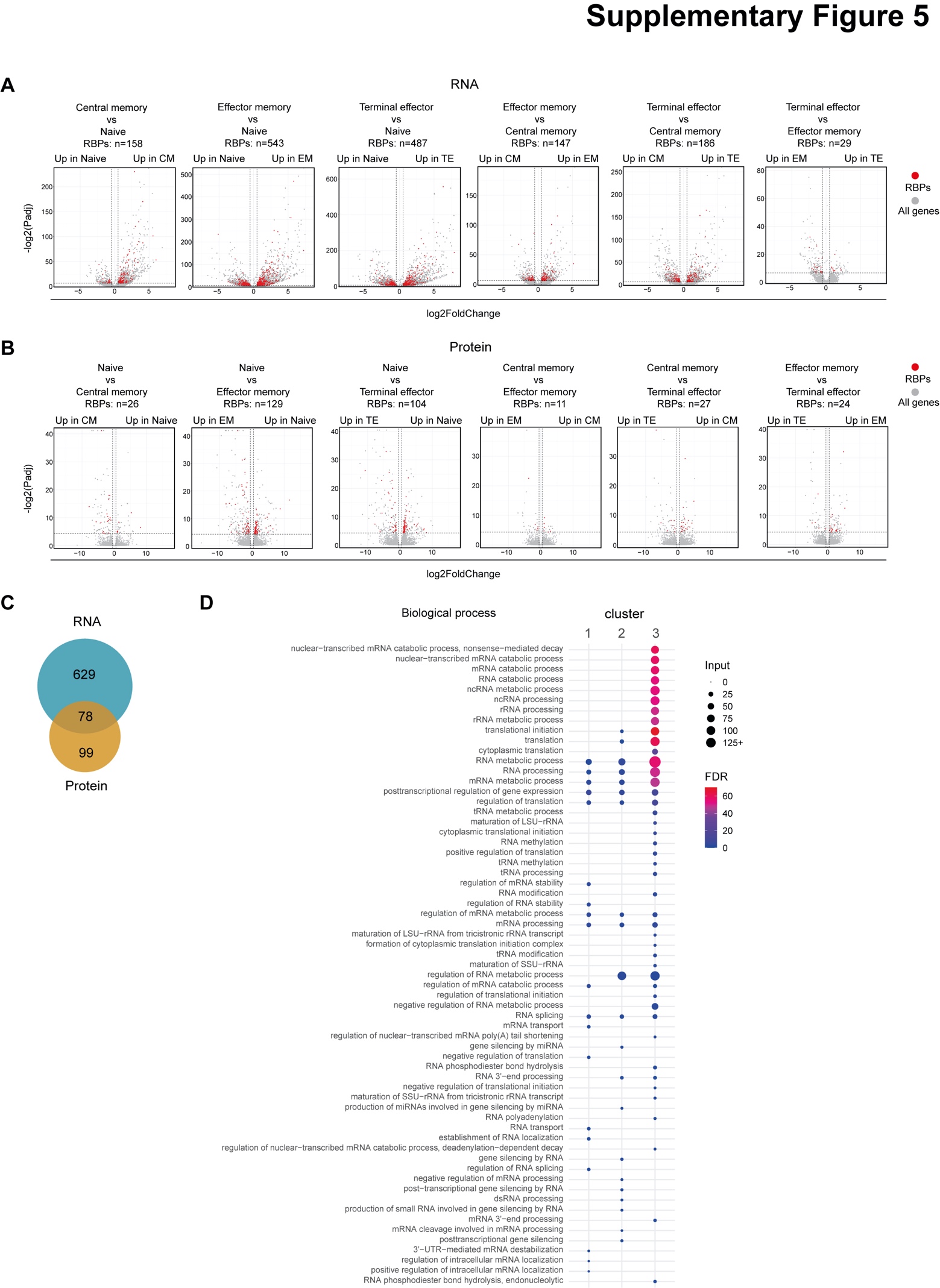

**Supplementary Figure 5:** **Differential RBP expression and GO-term analysis of RBPs in human CD8^+^ T cells.**

(**A-B**) Volcano plots of differential RNA **(A**) and protein expression **(B)** of all genes (gray) and of RBPs (red) between human CD8^+^ T cell subsets. (**C**) Venn diagram showing the overlay between RBP RNA and protein expression. (**D**) RNA-related biological processes enriched in clusters identified in Figure 5B (FDR<0.05).

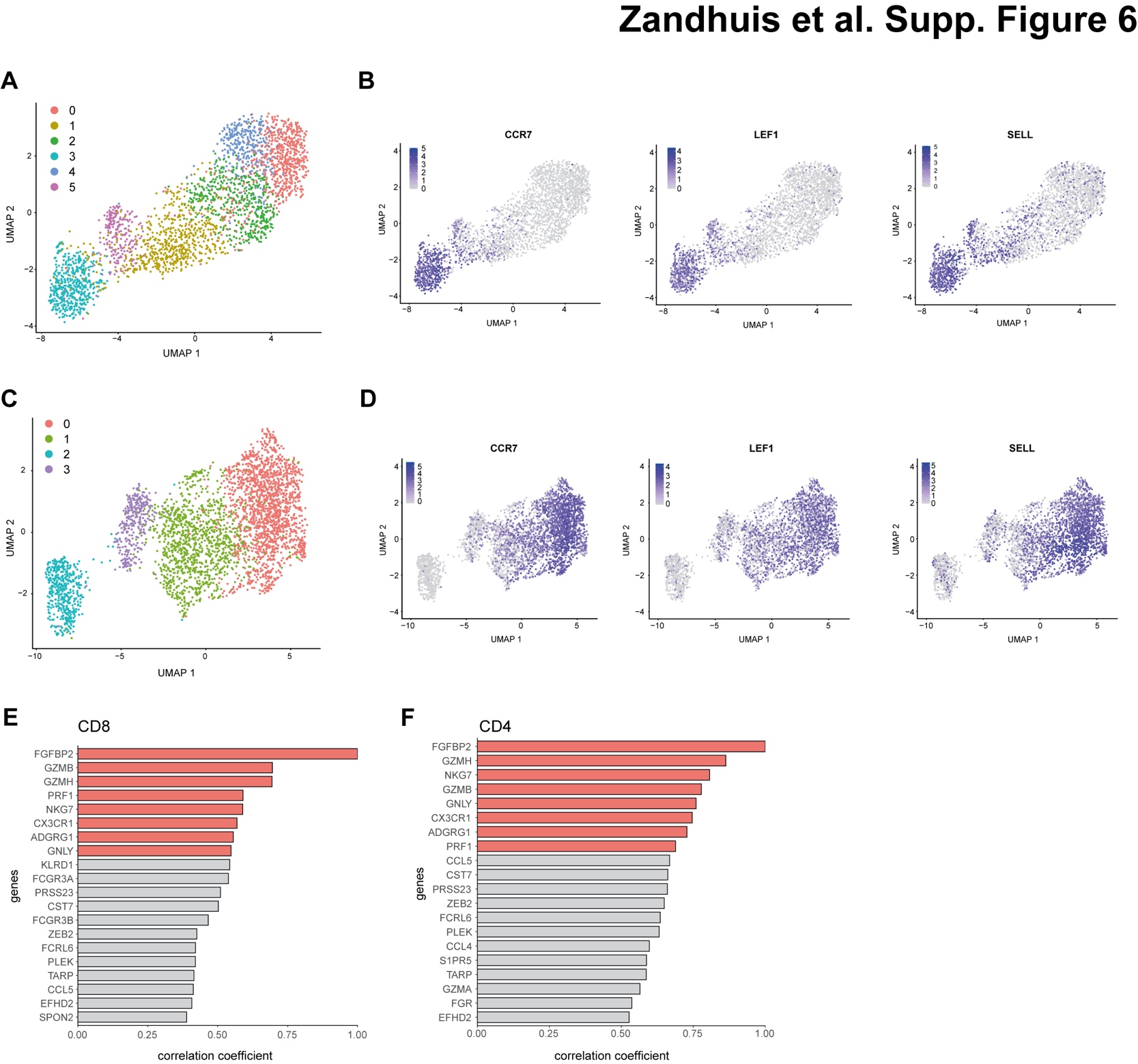

**Supplementary Figure 6:** **Identification of the cytotoxic gene signature in non-naïve CD4^+^ T cells and CD8^+^ T cells.**

(**A, C**) Uniform Manifold Approximation and Projection (UMAP) plot followed by unsupervised clustering on single-cell RNA sequencing data from blood-derived CD8^+^ T cells (A) and CD4^+^ T cells (C; *see Methods)*. (**B, D**) Expression levels of *CCR7*, *LEF1* and *SELL* among CD8^+^ T cells (B) and CD4^+^ T cells (D). (**E, F**) Bar chart depicting the 20 genes most correlated with *FGFBP2* across non-naïve CD8^+^ T cells (E), and non-naïve CD4^+^ T cells (F).

**Supplementary Table 1: RBD list and aggregated RBP reference list.**

**Supplementary Table 2: Subset specific expressed and differentially expressed RBPs between B cells, CD4 T cells and CD8 T cells.**

**Supplementary Table 3: Differentially expressed RBPs between CD19+ B cell subsets.**

**Supplementary Table 4: Differentially expressed RBPs between CD4+ T cells.**

**Supplementary Table 5: Differentially expressed RBPs between CD8+ T cells.**

**Supplementary Table 6: Differentially expressed RBPs between CD8+ T cells.**

**Supplementary Table 7: Enriched biological processes among RBP clusters identified in CD4+ T cells and CD8+ T cells.**
